# A DARPin-based molecular toolset to probe gephyrin and inhibitory synapse biology

**DOI:** 10.1101/2022.06.30.498253

**Authors:** Benjamin F. N. Campbell, Antje Dittmann, Birgit Dreier, Andreas Plückthun, Shiva K. Tyagarajan

## Abstract

Neuroscience currently requires the use of antibodies to study synaptic proteins, where antibody binding is used as a correlate to define the presence, plasticity, and regulation of synapses. Gephyrin is an inhibitory synaptic scaffolding protein used to mark GABAergic and glycinergic postsynaptic sites. Despite the importance of gephyrin in modulating inhibitory transmission, its study is currently limited by the tractability of available reagents. Designed Ankyrin Repeat Proteins (DARPins) are a class of synthetic protein binder derived from diverse libraries by *in vitro* selection, and tested by high-throughput screening to produce specific binders. In order to generate a functionally diverse toolset for studying inhibitory synapses, we screened a DARPin library against gephyrin mutants representing both phosphorylated and dephosphorylated states. We validated the robust use of anti-gephyrin DARPin clones for morphological identification of gephyrin clusters in rodent neuron culture and brain tissue, discovering previously overlooked clusters. This DARPin-based toolset includes clones with heterogenous gephyrin binding modes that allowed for identification of the most extensive gephyrin interactome to date, and defined novel classes of putative interactors, creating a framework for understanding gephyrin’s non-synaptic functions. This study demonstrates anti-gephyrin DARPins as a versatile platform for studying inhibitory synapses in an unprecedented manner.

## Introduction

Biological research has relied for decades on the accuracy and precision of specific antibodies to morphologically describe protein localization and dynamics, or to biochemically describe protein interaction partners, using techniques such as immuno-labelling, immunoprecipitation, and immunoassays, amongst others. While antibody-based tools have been invaluable, for a given protein we often lack a variety of binders which perform excellently across applications. Antibodies that detect fixed proteins in tissue (which are typically partially denatured), may not bind with the same affinity or specificity to the same protein in a lysate (which may retain a more native confirmation). The heterogeneous quality of some commercial antibodies presents an additional challenge as the often ambiguous or unknown antibody sequence, provenance, and specificity of poly- and monoclonal antibodies alike leads to false information and ultimately a high additional cost to research (Bradbury & Plückthun, 2015; “Protein Binder Woes,” 2015). This problem is especially relevant for the study of synaptic proteins, be they receptors or scaffolds, as these proteins are often used as markers to define the presence, plasticity, and regulation of synapses as a strong correlate for synaptic function. For example, ionotropic glutamate receptor subunits and the scaffolding molecule PSD-95 are frequently used to define the excitatory post-synapse, while GABA_A_ receptors (GABA_A_Rs) and the scaffolding protein gephyrin define the inhibitory post-synapse (Micheva et al., 2010).

Gephyrin is a highly conserved signaling scaffold which oligomerises into multimers and binds to cognate inhibitory synaptic proteins to functionally tether GABA_A_Rs at postsynaptic sites in apposition to pre-synaptic GABA release sites (Tyagarajan & Fritschy, 2014). Gephyrin is composed of 3 major domains: the N-terminal G domain and C-terminal E domain facilitate self-oligomerization of a gephyrin lattice underneath inhibitory postsynaptic sites, and they are linked together by the C domain which is a substrate for diverse posttranslational modifications (Sander et al., 2013; Tyagarajan & Fritschy, 2014). Gephyrin mediates its scaffolding role by coordinating the retention of inhibitory synaptic molecules (Fig 1A) including GABA_A_ and glycine receptors (GABA_A_Rs, GlyRs), collybistin, and neuroligin 2 through interactions at locations within the E domain or E/C domain interface (Choii & Ko, 2015; Tyagarajan & Fritschy, 2014), with additional protein interactors binding to the G and C domains. Therefore, via homo- and heterophilic protein-protein interactions, gephyrin can control inhibitory post-synaptic function.

**Figure 1.**
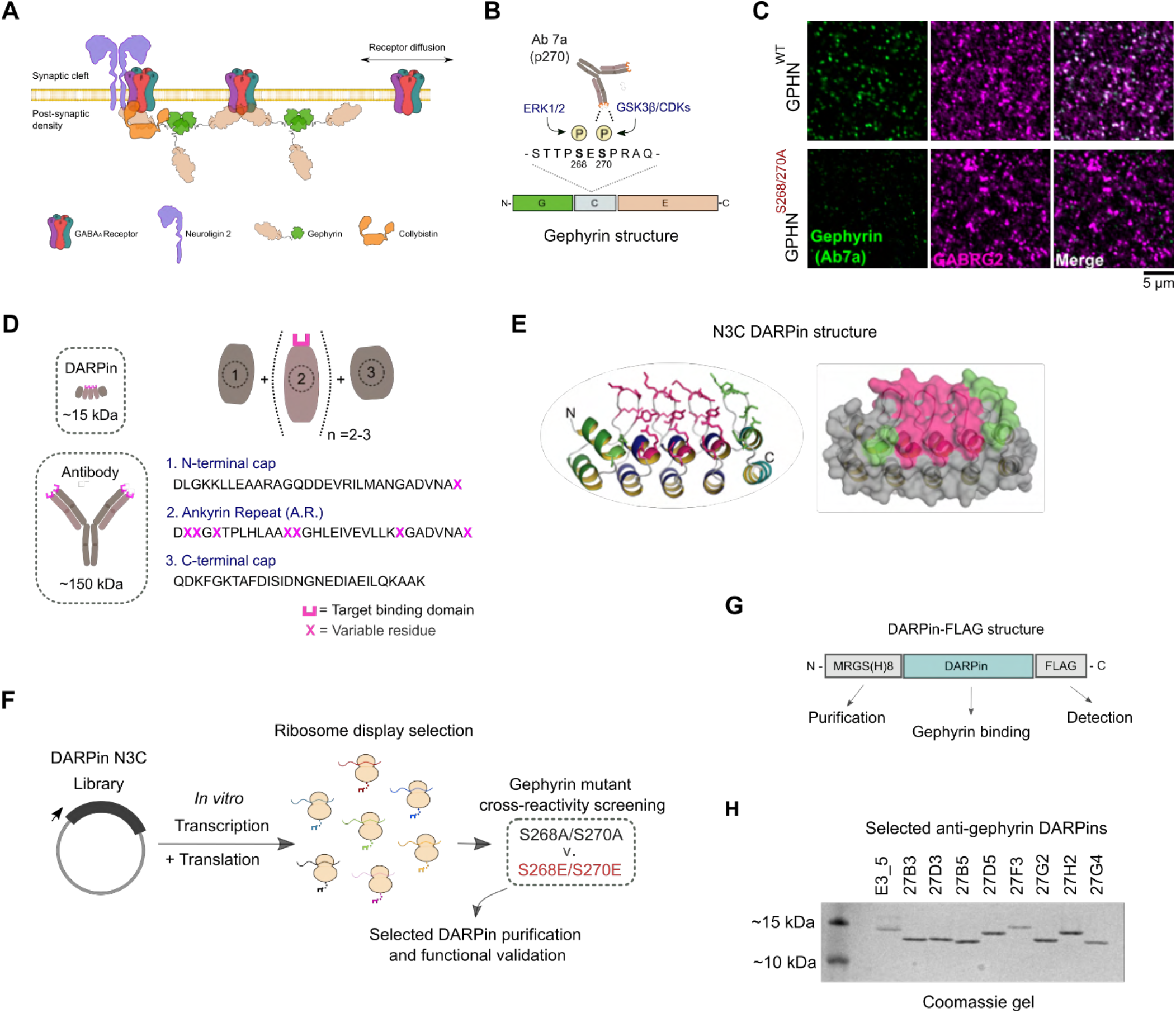
*In vitro* selection and generation of anti-gephyrin DARPins. **A)** Diagram of gephyrin function at the inhibitory post-synapse via its scaffolding role. **B)** Gephyrin domain structure and location of key phospho-serine residues S268 and S270, the commonly used antibody clone for detection of gephyrin (Ab7a) is pS270-specific. **C)** The antibody Ab7a does not detect gephyrin clusters colocalised with the γ2 GABA_A_ receptor subunit (GABRG2) in a phospho-null mouse model where S268 and S270 are mutated to alanines. **D)** DARPins are an order of magnitude smaller than conventional antibodies and achieve target binding specificity by varying the sequence of ankyrin repeats (A.R.) with variable residues (magenta). **E)** DARPin library design, with residues in magenta randomized in the original design and additional residues randomized in the caps (green). An N3C structure is shown with the N-cap as a green ribbon and the C-cap as a cyan ribbon with green side chains. **F)** Schematic anti-gephyrin DARPin selection and screening. **G)** Structure of DARPin-FLAG clones used for initial validation experiments contain an N-terminal His_8_ tag and C-terminal FLAG tag for purification and detection respectively. **H)** Coomassie-stained gel of the non-binding control (E3_5) and eight anti-gephyrin DARPin binders. **Figure 1 – Source data 1:** raw image and annotated uncropped Coomassie gel from Figure 1 H.

Gephyrin’s scaffolding role is dynamically regulated by its post-translational modifications (PTMs). Gephyrin phosphorylation at several defined serine residues controls gephyrin oligomerisation /compaction and thereby affect GABAergic transmission (Battaglia et al., 2018; Ghosh et al., 2016; Petrini & Barberis, 2014; Zacchi et al., 2014). Two of these phospho-sites, serines S268 and S270, are targeted by the kinases ERK1/2 and GSK3ß or cyclin-dependent kinases (CDKs) respectively, to downregulate gephyrin clustering (Fig 1B), thereby controlling post-synaptic strength (Tyagarajan et al., 2013). These phosphorylation events directly regulate gephyrin conformation via packing density changes to alter GABA_A_ receptor dwell time (Battaglia et al., 2018), by altering gephyrin interacting partners (Zhou et al., 2021), or some combination of the two (Specht, 2019). Unfortunately, the most widely used anti-gephyrin antibody for identifying inhibitory postsynaptic sites, monoclonal antibody clone Ab7a, is sensitive to phosphorylation at serine 270 (Kalbouneh et al., 2014; Kuhse et al., 2012; Zhou et al., 2021), thus complicating interpretation of inhibitory postsynaptic presence, size, or dynamics.

In addition to PTMs, gephyrin is regulated by alternative splicing by a suite of exonic splice cassette insertions (annotation outlined in (J.-M. Fritschy et al., 2008)). While the principal (P1) isoform of gephyrin in neurons facilitates its synaptic scaffolding role, gephyrin is also a metabolic enzyme which participates in molybdenum cofactor (MOCO) biosynthesis (Nawrotzki et al., 2012; Schwarz & Mendel, 2006; Tyagarajan & Fritschy, 2014). MOCO synthesis can be mediated in non-neuronal cells by an isoform that includes the C3 splice cassette (Licatalosi et al., 2008; Meier et al., 2000; Smolinsky et al., 2008), suggesting that gephyrin harbors both isoform- and cell-type-specific functions.

Gephyrin has been reported to complex with a wide variety of proteins as determined by both targeted and unbiased interaction studies (Fuhrmann et al., 2002; Sabatini et al., 1999; Uezu et al., 2016). These screens have implicated gephyrin in non-synaptic processes including regulation of mTOR signaling (Sabatini et al., 1999; Wuchter et al., 2012), and motor protein complexes (Fuhrmann et al., 2002). Furthermore these interactomes have identified novel proteins such as InSyn1, with implications in understanding the heterogeneity of inhibitory synapse organisation (Uezu et al., 2019). Still, the overlap in coverage of gephyrin’s interactome in each study has been variable with respect to identification of canonical inhibitory synaptic proteins due to limitations of each screening technique. Taken together, there is a need to generate and characterise molecular tools that can 1) interrogate gephyrin in different applications 2) be functionally validated for the experiment in question, and 3) be diverse enough in their mode of interaction to not limit the different protein functional states that can be probed.

Designed Ankyrin Repeat Proteins (DARPins) represent an attractive alternative tool compared to conventional antibodies as they are highly stable and specific synthetic protein binders, which can be generated via high-throughput *in vitro* selection and screening (Binz et al., 2004; Kohl et al., 2003). Since they possess a defined genetic sequence, they can be adapted into diverse fusion constructs, and their structural stability facilitates their engineering to achieve differential binding (Harmansa & Affolter, 2018; Plückthun, 2015). DARPins are composed of a variable number (typically 2-3) ankyrin repeats containing randomised residues, flanked by N- and C-terminal capping repeats with a hydrophilic surface that shield the hydrophobic core. Each repeat forms a structural unit, which consists of a β-turn followed by two antiparallel α-helices and a loop reaching the turn of the next repeat. The randomised residues on adjacent repeats within the β-turn turns and on the surface of the α-helices form a variable and contiguous concave surface that mediates specific interactions with target proteins. Using a DARPin library with high diversity (approx. 10^12^ unique DARPins), DARPins can be selected using ribosome display and then screened for particular binding characteristics (Dreier & Plückthun, 2012; Plückthun, 2012). Using this approach, DARPins have been shown to selectively bind to different conformations of proteins, include those brought about by phosphorylation (Kummer et al., 2012; Plückthun, 2015).

Despite being used extensively as both experimental tools for structural biology as well as therapeutics (Plückthun, 2015; Tamaskovic et al., 2012), DARPins have not yet been applied to neuroscience research in the current literature. In order to generate a new toolset of anti-gephyrin binders, we screened a DARPin library for binding to different gephyrin phosphorylation mutants and characterised the resulting DARPins in both morphological and biochemical applications. We validated the use of anti-gephyrin DARPins to understand how different binders can reveal novel aspects of gephyrin and inhibitory synapse biology highlighting heterogenaity of inhibitory post-synapse morphology and composition.

## Results

### Generation and selection of anti-gephyrin DARPins

Gephyrin clusters GABA_A_ receptors and other inhibitory molecules such as neuroligin 2 and collybistin at post-synaptic sites (Fig, 1A) where its clustering role is modified by phosphorylation, importantly at serines S268 and S270 (Fig. 1B). This phosphorylation of gephyrin links upstream signaling (e.g. neurotrophic factors, activity) to downstream gephyrin regulation of inhibitory synaptic function (Groeneweg et al., 2018; Tyagarajan & Fritschy, 2014). The commonly used commercial antibody clone for morphological detection of synaptic gephyrin (clone Ab7a) has been used extensively for almost four decades in the literature to identify inhibitory synapses (Pfeiffer et al., 1984). Though, rather than binding gephyrin regardless of its modified state, this antibody was recently demonstrated to specifically recognise gephyrin phosphorylated at serine S270 (Kuhse et al., 2012). This antibody’s specificity for phospho-gephyrin complicates interpretation of synaptic gephyrin cluster identification when using clone Ab7a, and prevents accurate detection of postsynaptic gephyrin clusters when gephyrin S270 phosphorylation is low or blocked. This is illustrated by the lack of binding of Ab7a to gephyrin in brain tissue derived from a phospho-S268A/S270A phospho-mutant mouse line, in which serines S268 and S270 are mutated to alanines (Fig 1C). Therefore, to generate protein binders that can more robustly identify gephyrin independently of its phosphorylation status, we looked beyond antibody-based binders to Designed Ankyrin Repeat Proteins (DARPins).

DARPins are small (∼12-15 kDa) compared to conventional antibodies (Fig. 1D), and their binding to specific target proteins is mediated by several randomised residues contained within assemblies of 2-3 variable ankyrin repeats (AR) flanked by capping repeats (Binz et al., 2004; Kohl et al., 2003). This basic DARPin structure creates a rigid concave shape with enhanced thermostability (Fig. 1E). In addition, DARPins do not contain cysteines, allowing for functional cytoplasmic recombinant expression in *E. coli* as well as cytoplasmic expression and functional studies in mammalian cells. We performed a ribosome-display selection, followed by screening of individual clones against recombinant gephyrin (P1 principal isoform) containing either S268A/S270A or S268E/S270E mutations (Fig. 1F) which mimic the respective de-phosphorylated and phosphorylated state, thus representing functionally distinct gephyrin conformations (Battaglia et al., 2018; Tyagarajan et al., 2013). This allowed us to define sensitivity towards the modified state and to widen the spectrum of DARPins obtained from the selection. Single DARPin clones were expressed in *E. coli* containing an N-terminal MRGS(H)_8_ (His_8_) tag and C-terminal FLAG tag (Fig. 1G). Initial screening was performed with 376 DARPin clones using a high-throughput HTRF assay with crude extracts derived from 96 well expression plates. Of the initial hits, 32 were sequenced and 25 unique DARPins identified. These DARPins were further screened using an ELISA-based assay for relative binding to the phospho-null or phospho-mimetic gephyrin isoforms, or the absence of target as control (Fig. 1 Suppl. 1). From this screen, eight DARPins were chosen for expression/purification and further analysis due to their high signal-to-background characteristics, as well as for equal binding to both phospho-mutant forms of gephyrin (Fig. 1 H, Fig. 1 Suppl. 1). These eight DARPins showed diversity in the variable residues in the target protein interaction surface, highlighting the broad spectrum of binders that were obtained with this technology, and suggesting that they likely interact with gephyrin using different binding orientation or epitopes and independent of phosphorylation (Fig. 1 Suppl. 2).

### Characterisation of anti-gephyrin DARPins as morphological tools

The antibody clone Ab7a has been used extensively to both define the location, size, and dynamics of postsynaptic gephyrin puncta (Bausen et al., 2010; Kalbouneh et al., 2014; Niwa et al., 2019). However, this antibody reacts preferentially with gephyrin phosphorylated at S270, and sometimes also labels non-specific structures such as the nucleus (Fig. 2A). Alternative anti-gephyrin antibodies exist such as clone 3B11 which can be used for immunoprecipitation of gephyrin and detection on immunoblots, but leads to high background when used to label synapses (Fig. 2 Suppl. 1B). To determine whether anti-gephyrin DARPins function as antibody-like tools in tissue staining (in addition to binding recombinant gephyrin *in vitro*), we compared FLAG-tagged anti-gephyrin DARPins against antibody clone Ab7a for staining in primary rat hippocampal neuron culture at 15 days *in vitro* (DIV) (Fig. 2 Suppl. 1. A-C). While the unselected control DARPin clone E3_5-FLAG (Binz et al., 2003) did not present with detectable signal (Fig. 2A), DARPin-FLAG clones 27B3, 27D3, 27F3, and 27G2 labelled gephyrin puncta with high specificity (Fig. 2 A, Suppl. Fig 2A, B, C). Clone 27D5-FLAG produced no detectable signal, and clones 27B5, 27H2, and 27G4 labelled gephyrin puncta but produced considerable background comparable to another commercial anti-gephyrin antibody (clone 3B11) (Suppl. Fig. 2B). Moreover, clones 27B3, 27D3, 27F3, and 27G2 colocalised with presynaptic vesicular GABA transporter (VGAT)-containing axon terminals (Fig. 2B). We compared the fraction of detected gephyrin puncta colocalised with VGAT, as well as the size of detected gephyrin clusters, using both the antibody Ab7a and selected DARPin-FLAG clones that displayed low background namely 27B3, 27D3, 27F3, and 27G2 (Fig. 2C, Fig. 2D). We found no differences between DARPin-FLAG 27B3 or 27G2 and Ab7a colocalisation with VGAT indicating equal functionality in morphological applications. DARPin-FLAG 27D3 and 27F3 labelled puncta of a smaller size, which could relate either to their affinity for synaptic gephyrin or heterogeneity in epitope accessibility as different postsynaptic gephyrin puncta may differ in their isoform or post-translationally modified state.

**Figure 2.**
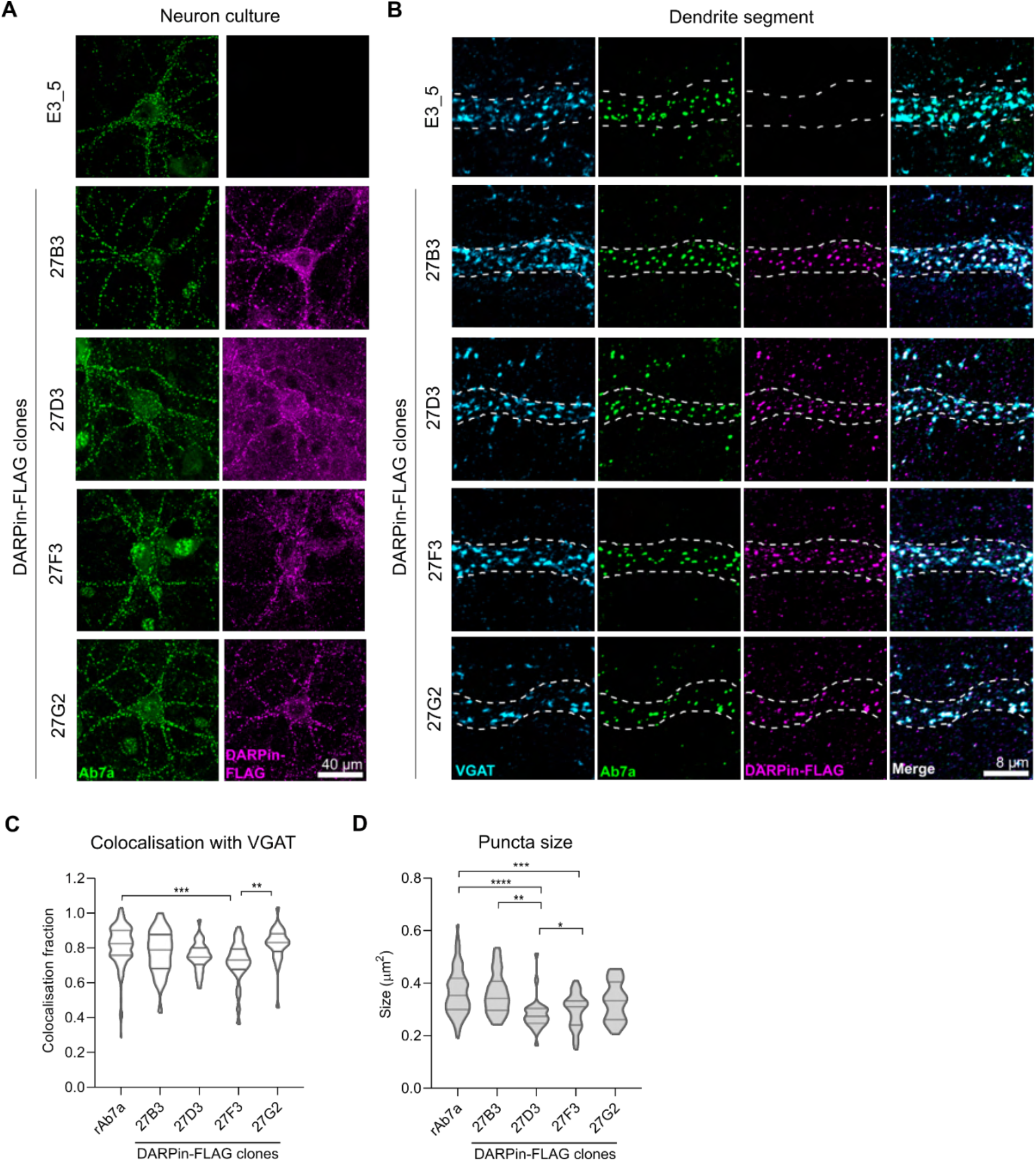
Anti-gephyrin DARPins specifically label gephyrin at inhibitory postsynaptic sites. Native gephyrin in fixed hippocampal neuron cultures (DIV15) probed using DARPin-FLAG clones, subsequently detected with anti-FLAG antibodies, and compared to staining with commercial anti-gephyrin antibody clone Ab7a. **A)** Representative images of DARPin-FLAG clones 27B3, 27D3, 27F3, and 27G2 gephyrin puncta colocalised to Ab7a signal compared to the control DARPin E3_5. **B**) Higher magnification images of dendrite segments showing detected DARPin-FLAG signal colocalised with pre-synaptic VGAT. **C**) Colocalisation analysis indicating the fraction of gephyrin puncta that colocalize with VGAT along a proximal dendrite segment (> 30 neurons/group pooled across 3 experiments)**. D)** Average puncta size identified by antibody Ab7a or DARPin-FLAG clones averaged by cell (pooled across neurons, >1100 synapses/group pooled across 3 experiment). **Statistics:** Panels C+D: One-way ANOVA, Tukey post-hoc test comparing all groups **** p<0.0001, *** p<0.0005, ** p<0.005 * p<0.05. Figure 2 – Source data 1. Contains the data and statistical analysis to generate the violin plot in panels C and D.

### Anti-gephyrin DARPin-hFc fusion construct identifies phosphorylated and non-phosphorylated gephyrin clusters in mouse brain tissue

Identification of inhibitory synapses often involves the co-labelling of both pre-and postsynaptic structures using multiple antibodies raised in different species. In order to label gephyrin clusters in the brain we replaced the His_8_ and FLAG epitope tags from DARPin-FLAG clones 27B3, 27F3, 27G2 and the control clone E3_5 with an N-terminal human serum albumin (HSA) leader sequence and C-terminal human Fc (hFc) tag for mammalian recombinant production and purification and detection (Fig. 3. Suppl. 1.). The addition of the hFc tag allows for use in tandem with essentially all primary antibodies targeting synaptic markers raised in commonly used species such as rat, mouse, rabbit, goat, and guinea pig. Furthermore, it makes the construct bivalent. Consistently, DARPin-hFc 27G2 specifically labeled gephyrin puncta apposed to presynaptic VGAT terminals in both hippocampal neuron culture and mouse brain tissue (Fig. 3. Suppl. 2.). The specificity of this labelling could be confirmed by incubating DARPin-hFc 27G2 with a molar excess of recombinant gephyrin as a competitor, which led to the loss of immunofluorescent signal (Fig. 3. Suppl. 3.).

A significant fraction of synaptic gephyrin clusters are phosphorylated at serine 270, and therefore lead to an uncertain interpretation when their size and dynamics are assessed using the phospho-specific antibody Ab7a (Kalbouneh et al., 2014; Specht, 2019; Zhou et al., 2021). As predicted, DARPins-hFc 27G2 can label gephyrin puncta in both wild-type and phospho-S268A/S270A mutant mouse tissue while the commercial pS270-specific antibody Ab7a does not (Fig. 3 A).

**Figure 3:**
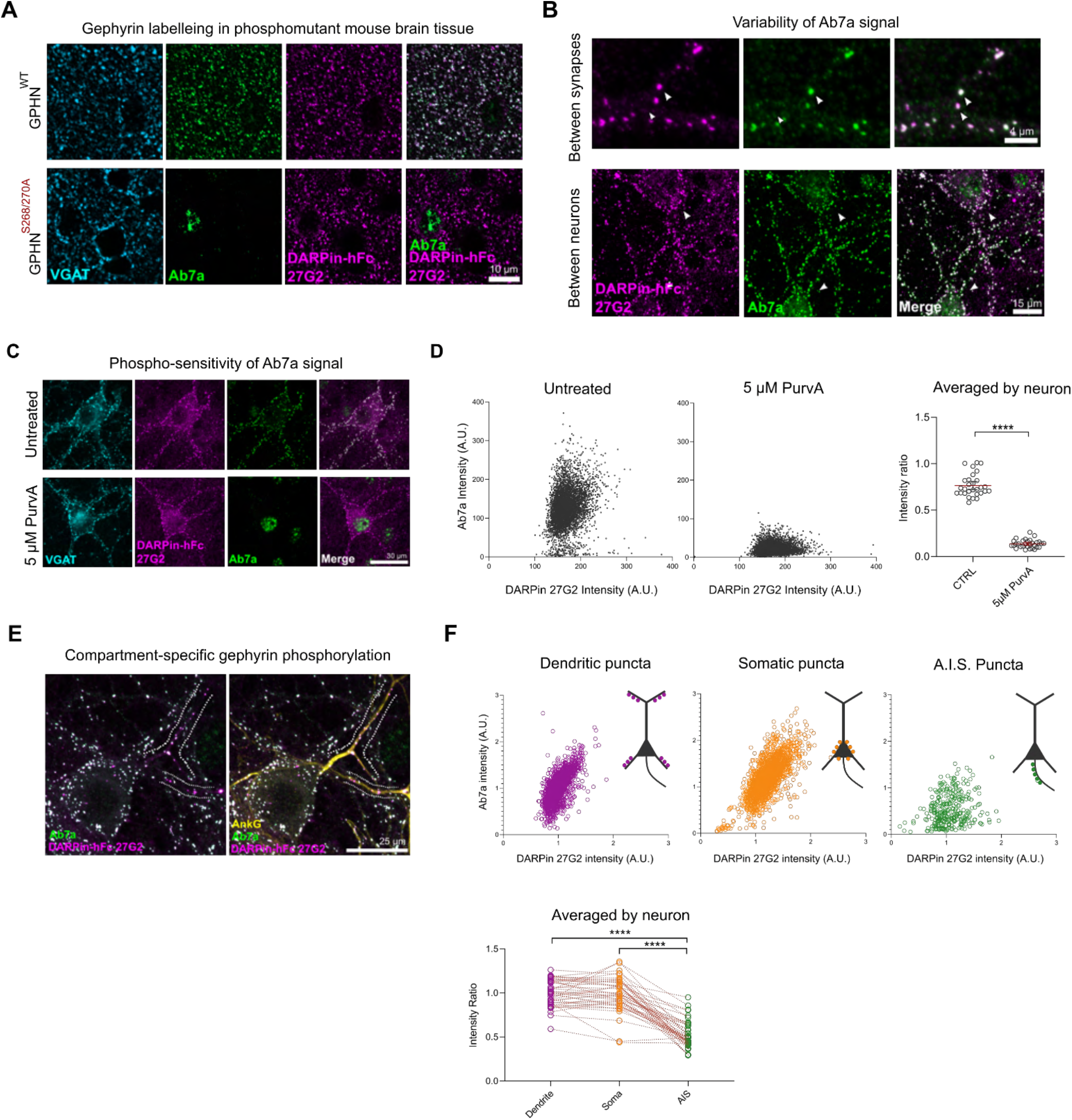
Phospho-insensitive DARPin-hFc 27G2 multiplexed with antibody Ab7a can assess synapse-specific gephyrin S270 phosphorylation. **A)** Representative images of DARPin-hFc 27G2 (but not antibody Ab7a) labelling gephyrin puncta in both wild-type (WT) and phospho-mutant gephyrin S268A/S270A mutant mouse brain tissue (somatosensory cortex layer 2/3). **B)** Representative images from hippocampal neuron culture showing the relative Ab7a signal (indicating S270 phosphorylation) varies by synapse and between neurons. **C)** Representative image showing DARPin-hFc 27G2 binding at synaptic puncta in primary hippocampal neuron culture is preserved after inhibition of CDKs following 24-hour treatment with 5 µM Aminopurvalanol (PurvA) while Ab7a staining is severely reduced. **D)** The relative fluorescence intensity at individual synapses (pooled from 30 neurons per group) showing a pronounced decrease in the average Ab7a/DARPin-hFc 27G2 intensity ratio. Quantification of Ab7a/DARPin-hFc 27G2 fluorescence signal averaged across cells pooled from 3 independent experiments, n=30 cells/group. **E)** Representative images of hippocampal neuron culture used for quantification of relative Ab7a/DARPin-hFc labelling of clusters on the soma, proximal dendrites, or the A.I.S. (AnkG). **F)** Ab7a/DARPin intensity ratio of individual synapses pooled from 45 cells over 3 independent experiments showing a decrease in A.I.S. cluster Ab7a staining. Lower: Quantification indicates significantly reduced A.I.S. Ab7a labelling of clusters compared to dendritic or somatic compartments. **Statistics:** Panels D: One-way ANOVA, Panel F: Repeated measures One-way ANOVA. All panels: * p<0.05, ** p<0.01, *** p<0.001, **** p<0.0001. Mean and SD are presented. Figure 3 – Source Data 1: contains values and statistical results used to generate panels D and F.

The relative amount of Ab7a to anti-gephyrin DARPin signal could be used as a proxy to estimate relative gephyrin S270 phosphorylation at synapses. Indeed, we found that the Ab7a signal varied considerably both between adjacent synapses within a neuron and between neurons (Fig 3B, Fig. 3. Suppl. 4.). We confirmed the phospho-sensitivity of this analysis method by inhibiting cyclin-dependent kinases (upstream of gephyrin S270 phosphorylation) using 5 µM Aminopurvalanol A applied for 24 hours. This treatment reduced Ab7a but not DARPin-hFc 27G2 signal as indicated by the decrease in the ratio between these two intensities seen both for individual synapses and when averaged by neuron (Fig. 3C, D). We therefore examined the Ab7a / DARPin-hFc 27G2 intensity ratio between the somatic, dendritic, and axon-initial segment (A.I.S.) compartments in primary hippocampal neuron culture (Fig 3 E, F), finding a significant reduction in Ab7a signal within the A.I.S. as defined by AnkyrinG immunolabelling (AnkG). Our results demonstrate that gephyrin phospho-S270 status varies between two neighbouring clusters within a dendrite segment and also for the first time we can label gephyrin within the A.I.S.

### DARPin-hFc 27G2 detects previously overlooked gephyrin clusters in brain tissue

Antibody-based identification of gephyrin clusters in the brain is widely used to identify inhibitory synaptic sites, but current reagents may only capture a subset of synaptic gephyrin clusters, namely those with gephyrin significantly phosphorylated at S270. Therefore, we extended our analysis of post-synaptic gephyrin clusters using DARPin-hFc 27G2 and the phospho-S270 specific antibody Ab7a to mouse brain tissue, using the hippocampal CA1 area as a model. The hippocampus is organised in a layered structure, stratifying somatic from dendritic compartments, with compartment-specific GABAergic interneuron innervation patterns well described (Pelkey et al., 2017). We found lamina-specific variability in relative gephyrin phosphorylation at S270, which was significantly elevated in the stratum oriens and stratum lacunosum moleculare compared to other layers (stratum pyramidale and radiatum) (Fig. 4 A, B, C). Within the stratum pyramidale, we noticed a population of large, relatively hypo-phosphorylated clusters (Fig. 4D, Fig. 4 Suppl. 1) reminiscent of axon initial segment (A.I.S.) synapses (Fig. 4E). Indeed, while DARPin-hFc 27G2 labels large gephyrin clusters apposed to presynaptic VGAT terminals, Ab7a reactivity within the A.I.S. is relatively weak (Fig. 4F). These hypo-phosphorylated clusters co-localise with the α2 GABA_A_ receptor subunit thought to be enriched at the A.I.S. (Lorenz-Guertin & Jacob, 2018), and span the length of the A.I.S. as defined by AnkG. Therefore, DARPin-hFc 27G2 can better assess postsynaptic gephyrin at the A.I.S. and at synapses where gephyrin phosphorylation is low. These data indicate that gephyrin clusters on the A.I.S. have likely gone un- or under-reported in the literature, which is meaningful when considering that threshold-based detection of gephyrin is used as a proxy for inhibitory synapse presence and function (Micheva et al., 2010; Schneider Gasser et al., 2006).

**Figure 4.**
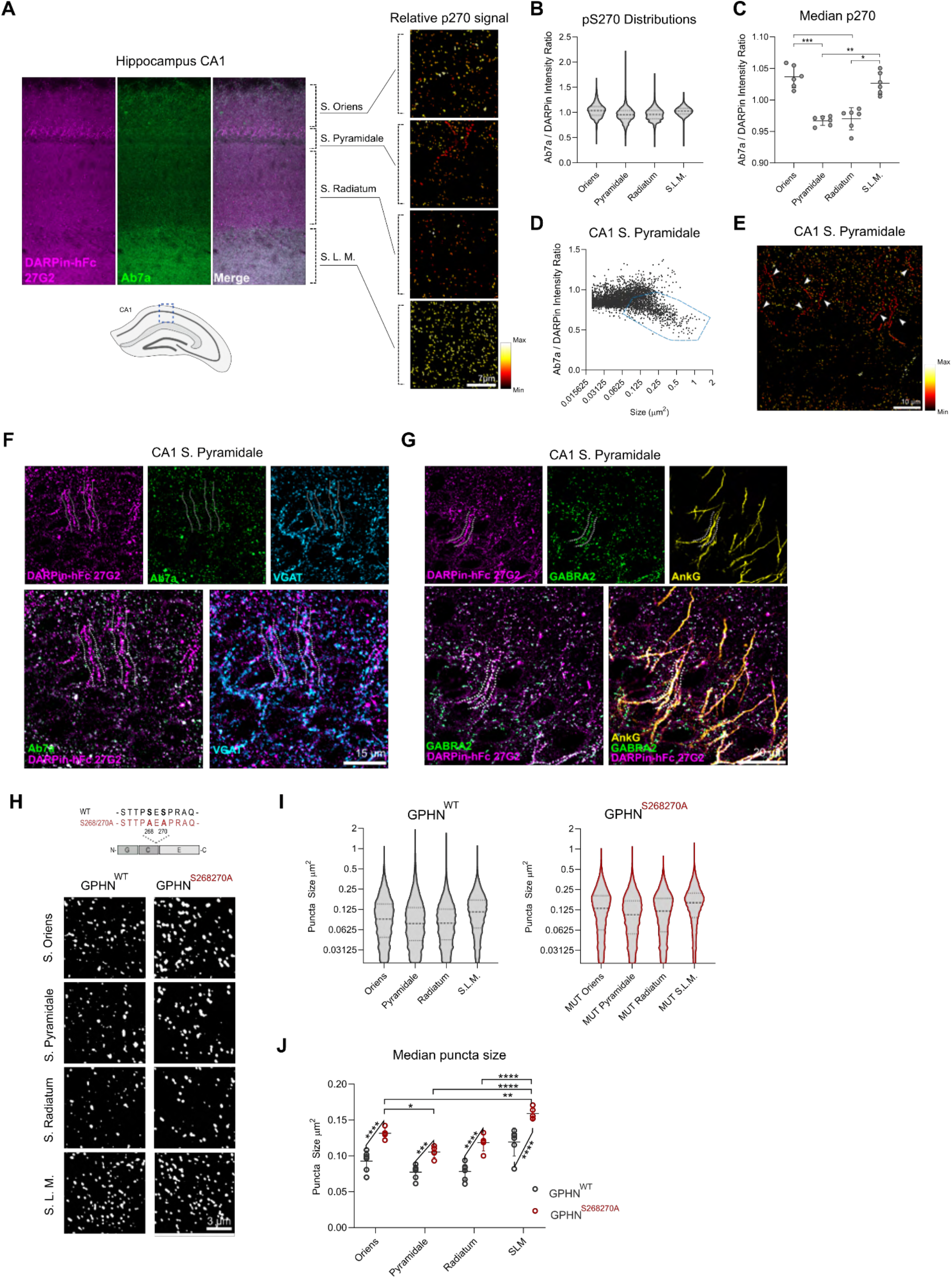
DARPin-hFc 27G2 labelling of gephyrin clusters demonstrates laminar and A.I.S.-specific S270 phosphorylation and phosphorylation-dependent cluster size regulation. **A**) Left: the relative Ab7a to DARPin-hFc 27G2 fluorescence intensity in the mouse hippocampus area CA1 shows layer-specific variability. Right: colourised gephyrin puncta indicating relative S270 phosphorylation as seen from hotter (more red/yellow) colouration. **B**) Distribution of relative gephyrin phosphorylated at S270 (p270) at puncta between hippocampal lamina. Data pooled between 6 adult mice, 3 sections analysed per mouse encompassing 14,000-47,000 gephyrin puncta per layer. **C)** Analysis of the median relative gephyrin pS270 ratio between hippocampal layers (data pooled between sections per mouse, n=6 mice quantified). **D)** Example distribution of gephyrin pS270 signal by puncta size in the CA1 stratum pyramidale, with a population of large, hypo-phosphorylated clusters outlined. **E)** Representative image of s. pyramidale with hot colours indicating gephyrin clusters with elevated phosphorylation, arrows indicate trains of large hypo-phosphorylated clusters. **F)** Representative image showing large DARPin-identified gephyrin clusters apposed to presynaptic VGAT-containing terminals with corresponding low Ab7a antibody signal. **G)** Representative image indicating gephyrin clusters on the A.I.S. (AnkG) colocalise with the α_2_ GABA_A_ receptor subunit. **H)** Representative images of gephyrin puncta identified using cluster analysis software in WT and S268A/S270A phospho-null mutant mice in the hippocampus using identical imaging parameters. **I)** Violin plots indicating the distribution of gephyrin puncta sizes (14,000-47,000 puncta per group, pooled across 5-6 mice per group). **J)** Analysis of the median puncta size between hippocampal layers and genotypes indicating larger gephyrin clusters in mutant mice. **Statistics:** Panels C: One-way ANOVA, Panel J: Mixed effects analysis comparing hippocampal lamina (horizontal bars) and genotypes (angled bars). All panels: * p<0.05, ** p<0.01, *** p<0.001, **** p<0.0001. Median and SD are presented. Figure 4 – Source Data 1: Contains data and statistical analysis presented in Figure 4 panels B, C, D, I, and J.

While gephyrin phosphorylation at S268 and S270 is thought to reduce gephyrin cluster size (Tyagarajan et al., 2013), the phospho-sensitivity of clone Ab7a has prevented our analysis of this relationship as this antibody does not react with dephosphorylated gephyrin (blocked in the mutant mouse). Therefore, we applied DARPin-hFc 27G2 to analyse gephyrin clusters in both WT and our phospho-null S268A/S270A mutant mouse model (GPHN^S268A/S270A^) (Fig 4H, I). We found that the median gephyrin cluster size is highest in the stratum oriens and stratum lacunosum moleculare in both WT and mutant mice, but that the median gephyrin cluster size is significantly enhanced across all layers when gephyrin phosphorylation is constitutively blocked in the S268A/S270A mutant mice (Fig. 4J). This represents the first confirmation that native gephyrin clusters in the brain are importantly regulated by serine 268 and 270 phosphorylation. Moreover, the identification of layer- and compartment-specific gephyrin phosphorylation in the hippocampus indicates that the use of DARPin-hFc binders may be a more robust morphological tool to investigate the heterogeneity of gephyrin and inhibitory synapses in the brain.

### Multiple gephyrin protein complex precipitations using unique DARPin binders establishes a consensus gephyrin interactome

Beyond applications for morphological detection of proteins in tissue, antibodies are essential for isolation of target protein complexes to understand their functional interaction networks. However, a network discovered by one binder may be different from another binder either due to affinity, or epitope accessibility involving targets in specific functional states. Gephyrin was first identified as a scaffolding protein, and yet throughout the past decades has been implicated additionally in complex signaling processes mediated by changes in its ability to interact with different protein partners. To gain a more complete picture of gephyrin binding partners, we precipitated native gephyrin protein complexes from mouse brain lysates with the traditionally used antibody clone 3B11 (suitable for immunoprecipitations) and each one of our DARPin-hFc clones 27B3, 27F3, 27G2, and the control DARPin E3_5 (Fig. 5 Suppl. 1). We then subjected the precipitated gephyrin complexes to interactor identification using quantitative liquid chromatography tandem mass spectrometry (LC-MS/MS) and compared the resulting interactomes (Fig. 5A, B). We considered proteins to be present when they were detected using at least two peptide signatures. Furthermore, we considered proteins as part of gephyrin complexes when they were present either only in the binder condition, or at least a log_2_ > 2.5-fold enriched in the binder condition over the control DARPin E3_5 with a false discovery rate (FDR)-adjusted p-value cut-off under 0.05 (Fig. 5B). These thresholds allow for a wider coverage to encompass most known interactors (Fig. 5 Suppl. 2) such as collybistin (ARHG9), GABA receptor subunits (GBRA1, 2), and a list of gephyrin interactors identified via BioID labelling (Uezu et al., 2016). Our results demonstrated that the abundance of canonical interactors spanned several orders of magnitude (Fig. 5 Suppl. 2) and provided enhanced coverage compared to the previously established BioID-determined interactome (Fig. 5C, Fig. 5 Suppl. 3). Each interactome differed by the number of identified proteins (Fig. 5C) where DARPin-hFc clones 27B3 and 27G2 identified 2-4 times more interactors than DARPin-hFc 27F3 or antibody 3B11, thus confirming the limitations of using only one binder to explore interacting protein networks.

**Figure 5:**
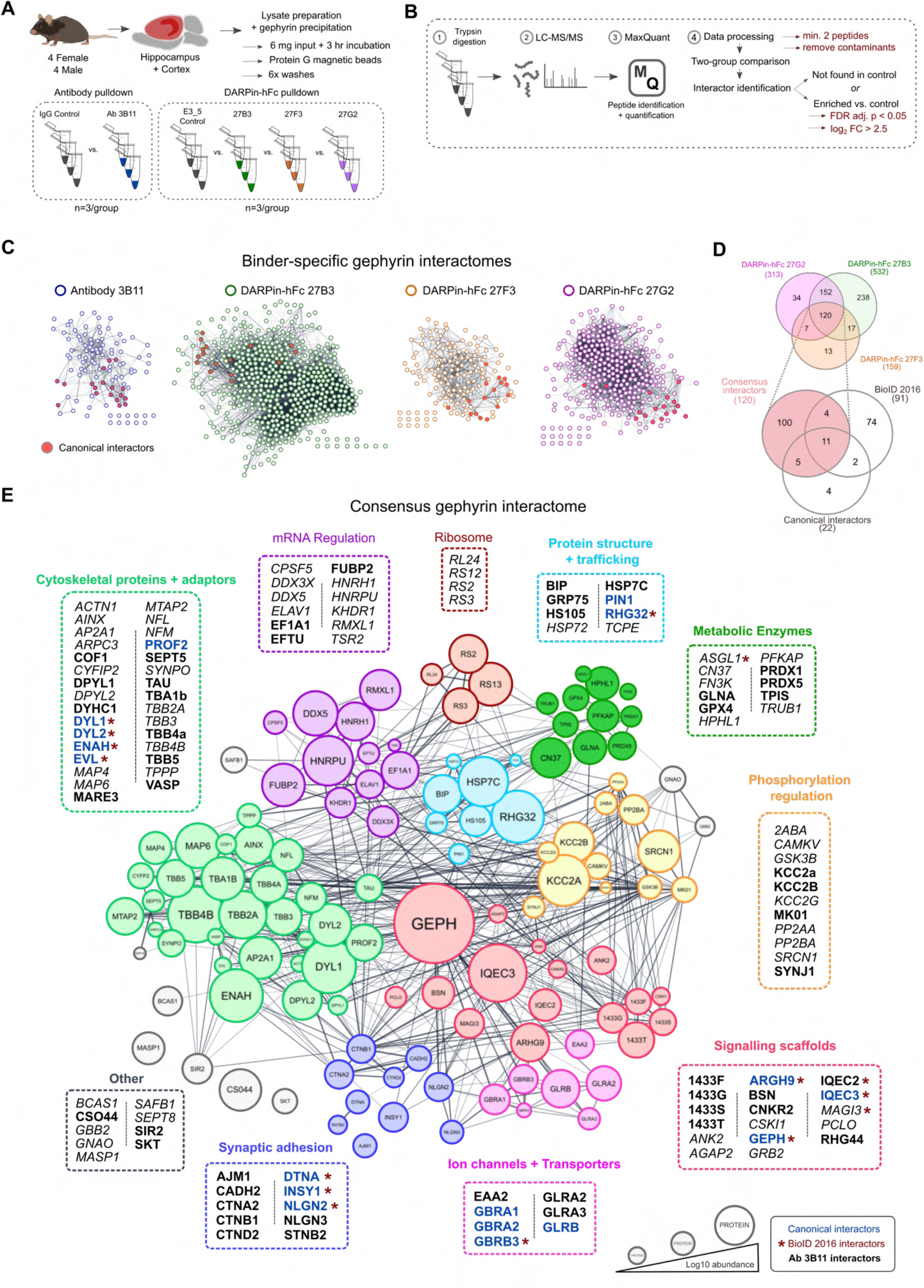
A DARPin-based consensus gephyrin interactome captures both known and novel protein interactors. **A**) Mouse brain tissue lysate preparation diagram. **B)** LC-MS/MS and interactome determination methodology workflow indicating thresholds for consideration of interacting proteins. **C)** Scale-free interaction networks (STRING) of gephyrin interactors identified from pulldowns using the commercial antibody 3B11, or DARPin-hFc 27B3, 27F3, and 27G2 compared to control conditions (containing antibody control IgG or the control DARPin-hFc E3_5). Nodes represent unique gephyrin interactors -red nodes indicate known (canonical) gephyrin interactors. **D)** Venn diagram of the overlap in identified interactors from gephyrin complexes isolated using different DARPin-hFc clones, bottom indicates coverage compared to an extensive gephyrin interactome determined using BioID labeling (Uezu et al., 2016) and 22 canonical gephyrin interactors identified from the literature. **E)** Consensus interactome of proteins identified by all DARPin-hFc clones and coloured by protein ontology. Canonical gephyrin interacting proteins are indicated by blue font, bold font indicates interactors also identified by the antibody clone 3B11. Asterisks indicate proteins previously identified by BioID (Uezu et al., 2016). Italic font indicates interactors exclusively identified by DARPins. Edges connecting protein nodes indicate putative interactions (STRING analysis), node circle size indicates relative protein abundance averaged across all experiments. Figure 5 – Source Data 1: List of interactors and relative abundance of detected proteins used to construct interaction networks and Venn diagrams in Figure 5 panels C, D, and E.

High-confidence interactome determination is limited both by the sensitivity of interactor detection as well as by the presence of false positives. Therefore, to compile a higher-confidence list of gephyrin interactors, we combined coverage between experiments using each DARPin-hFc clone to create a common gephyrin interaction network. We additionally cross-referenced this list with interactors precipitated by the antibody 3B11 as well as known binders identified from the literature to compile a high-confidence consensus gephyrin interactome (Fig. 5D), representing the largest compilation of putative gephyrin interactors to date. This network encompasses the majority of canonical gephyrin-associated proteins including GABA_A_ and glycine receptors, inhibitory synaptic scaffolding and adhesion molecules, and cytoskeletal adaptor proteins. As expected, over-representation analysis of the consensus interactome found significant enrichment for synaptic organization processes, but also unexpectedly those involved in protein trafficking, mRNA regulation, and metabolic processes (Fig. 5 Suppl. 4). Cataloguing of individual proteins by functional ontology revealed clusters of gephyrin interactors in mRNA regulation, cytoskeletal proteins and adaptors, metabolic enzymes and ribosomal subunits, together hinting at novel functions of gephyrin beyond synaptic scaffolding and molybdenum co-factor (MOCO) biosynthesis (Fig. 5E).

### Unique DARPin-hFc clones capture overlapping but ontologically distinct gephyrin interactomes

While our consensus gephyrin interactome may provide a robust framework to explore the related function of novel interacting proteins, the different scale of each network in terms of unique proteins identified and their different abundances suggests that each DARPin-hFc clone captures overlapping but unique gephyrin protein networks. To explore the extent of this phenomenon, we compared the relative abundance of interacting proteins which were constitutively present in all DARPin-hFc-derived gephyrin interactomes, and identified a subset of proteins, which showed significant variation in the abundance between the three DARPin-hFc-based pulldowns (Fig. 6 Suppl. 1). These included several canonical gephyrin interactors (Fig 6 A). For example, clone 27F3 precipitated significantly more IQEC3 (a guanine nucleotide exchange factor important for synapse specification (Früh et al., 2018)), while clone 27G2 captured gephyrin complexes containing more collybistin (ARHG9) (Fig. 6A). Binder-specific protein abundance profiles were more pronounced when examining non-canonical gephyrin interactor sets such as metabolic enzymes, mRNA binding proteins, and ribosomal subunits. These ontology groups demonstrated a consistently higher abundance in clone 27B3 and 27G2 compared to 27F3-based gephyrin interactomes. This differential interactor abundance could be due either to DARPins interacting with functionally distinct isoforms of gephyrin, or DARPin-specific interference with gephyrin conformation or interacting protein binding.

**Figure 6:**
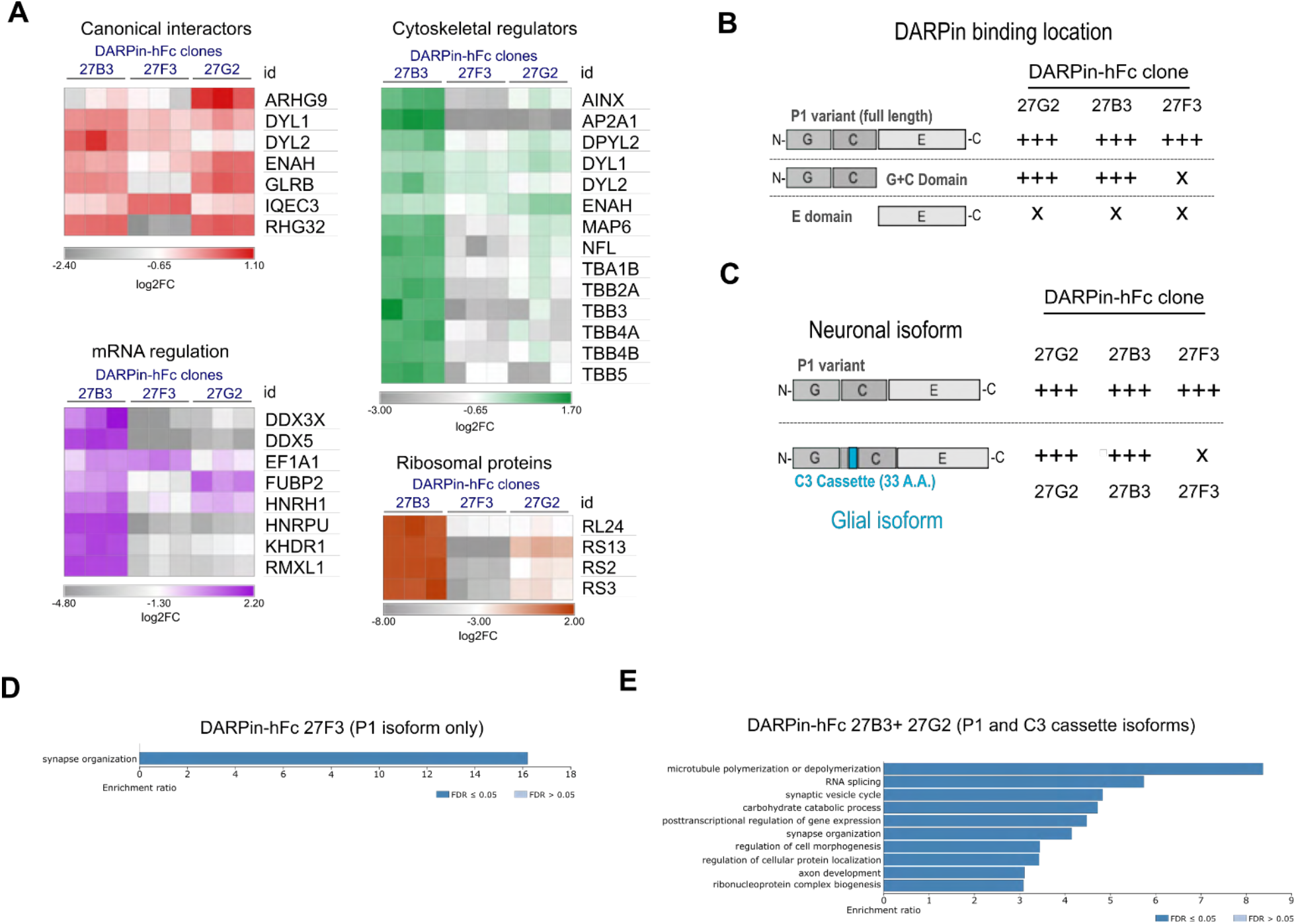
Diversity in DARPin-hFc clone-specific interactomes reveal putative isoform-specific gephyrin interactors. **A)** Canonical and non-canonical (metabolic, mRNA binding, and ribosomal ontology) gephyrin interactors show binder-specific abundance profiles. Only significantly regulated interactors are shown. **B)** DARPin-hFc clones 27B3 and 27G2 recognise both full length gephyrin and the GC-domain while clone 27F3 recognises only full length gephyrin suggesting different binding epitopes. **C)** DARPin-hFc 27F3 only recognizes the principal P1 (synaptic) isoform of gephyrin while clones 27B3 and G2 additionally recognize non-neuronal isoforms containing the C3 cassette. **D)** DARPin-hFc 27F3-determined gephyrin interactome enriched over-representation analysis of biological processes. **E)** DARPin-hFc 27B3 and 27G2-determined gephyrin interactome enriched over-representation analysis of biological processes. **Statistics:** Panel A: Two-way ANOVA with multiple comparisons correction comparisons all groups, 3 replicates per group. Figure 6 – Source Data 1: Values used to generate heat maps in Figure 6 panel A.

Gephyrin function is executed by several functional domains (G, C, and E domains), but it is also highly modified by phosphorylation as well as splice cassette insertions. To determine whether DARPin-hFc clones bind to different gephyrin domains or modified isoforms with different strength, we used an in-cell binding assay (Fig. 6 Suppl. 2) to assess the relative binding of these clones to different forms of eGFP-tagged gephyrin. As expected from the *in vitro* characterisation, there was no preference for any of the DARPin-hFc clones between wild-type gephyrin and the phospho-null or phospho-mimetic mutation-containing gephyrin at serines 268 and 270. Interestingly, we saw clear domain-specific binding preferences, with clones 27B3 and 27G2 interacting both with full-length gephyrin or the G and C domains in isolation, whereas clone 27F3 could only bind to full-length gephyrin (Fig. 6 B, Fig. 6 Suppl. 2). Gephyrin splice cassette C3 is constitutively spliced out in neurons by the splicing factor NOVA (Licatalosi et al., 2008), implying it is not needed for synaptic scaffolding. However, the C3 cassette is included in gephyrin expressed within non-neuronal cells where is contributes towards molybdenum cofactor (MOCO) synthesis activity (Smolinsky et al., 2008), or possible other functions (Fig. 6C). We found that the C3 cassette is significantly less detected by DARPin-hFc 27F3, while clones 27B3 and 27G2 bind to both the principal (P1) and C3-containing cassette isoforms equally (Fig. 6C, Fig. 6. Suppl. 2). We additionally probed for binding to gephyrin containing the C4a cassette (thought to be brain-enriched but without a clearly identified function). None of DARPin-hFc clones tested interacted strongly with the C4a-gephyrin isoform, while the antibody clone 3B11 interacted with this isoform at similar levels to the other gephyrin isoforms.

To understand whether the different DARPin-hFc clones can interact with ontologically distinct gephyrin protein networks, we performed over-representation analysis of proteins which are exclusive or significantly elevated in the interactome detected by clone 27F3 (neuronal isoform specific) or detected exclusively or significantly elevated by clones 27B3 and 27G2 (bind to neuronal and glial gephyrin isoforms). While we only saw enrichment for synaptic organization-related biological processes from DARPin-hFc 27F3 enriched interactors, we additionally found enrichment for cytoskeletal processes, ribosomal complex formation, and proteins involved in mRNA splicing and transport for the 27B3 and 27G2 enriched interactomes (Fig. 6 D, E). This suggests that the non-neuronal isoforms of gephyrin could be involved in these other distinct biological processes. In support of this hypothesis, when examining for proteins of glial or myelin ontology, we saw overall higher presence and abundance in the interactomes determined using clones 27B3 and 27G2 (Suppl. Fig. 6F). These data indicate that understanding the isoform-specificity of different DARPin clones will be useful for future dissection of gephyrin functionality at synapses, but also outside of synaptic sites or in non-neuronal cells.

## Discussion

In this study, we generated and characterised anti-gephyrin DARPins as a novel tools to study inhibitory synapse biology. This novel class of gephyrin protein binders specifically interacts with gephyrin in both morphological and biochemical applications to allow us to label gephyrin clusters and isolate gephyrin protein complexes without the limitations of previous antibody-based tools. We furthermore demonstrated that these DARPins can capture a greater diversity of gephyrin forms and functions, which will allow researchers to further characterise gephyrin and inhibitory synapses alike.

### Use of anti-gephyrin DARPins as morphological tools

Gephyrin is most widely used as an inhibitory postsynaptic marker due to its specific enrichment at inhibitory postsynaptic sites, but current antibody epitope limitations mask the heterogeneity of postsynaptic gephyrin clusters which can be probed. As our DARPins are insensitive to modification at two key phospho-sites thought to be dynamically regulated at synapses, we were able to identify previously masked gephyrin clusters at the axon initial segment where relative gephyrin S270 phosphorylation is low (and thus difficult to detect with the antibody Ab7a). Because most image analysis methods use threshold-based detection of gephyrin cluster presence and dynamics, A.I.S. gephyrin clusters (and identification of inhibitory synapses) will be massively underrepresented in the literature. For example, by using only the antibody Ab7a, gephyrin was suggested to play a less important role in scaffolding A.I.S. synapses (Gao & Heldt, 2016), whereas the large gephyrin clusters illuminated using DARPins suggests the opposite. Inhibitory input onto the A.I.S. provided by Chandelier interneurons plays an important role in gating neuronal output (Pelkey et al., 2017). Therefore, studying gephyrin A.I.S. dynamics is especially relevant for uncovering mechanisms of network plasticity and how inhibition controls circuit function. Outside of the A.I.S., we documented clear changes in relative gephyrin S270 phosphorylation in the stratum oriens and stratum lacunosum moleculare, indicating potential interneuron-specific or input-layer specific regulation of gephyrin function. Therefore, these DARPin-based tools can be used not only to robustly describe native gephyrin clusters in culture systems and in tissue, but they can also be used in tandem with gephyrin phospho-specific antibodies such as clone Ab7a to examine how genetics, environmental factors, or network activity regulate inhibitory adaptations via gephyrin. Moreover, DARPin binders may be able to better capture the heterogeneity of inhibitory postsynaptic sites that display differences in molecular composition regulation dependent on presynaptic inhibitory input (Chiu et al., 2018). The inclusion of the hFc tag on the DARPin constructs additionally allows them to be used with anti-human secondary reagents, and thus in conjunction with the vast majority of commercial and homemade antibodies against other synaptic markers raised in non-human species.

DARPins lack cysteines, and thus have an advantage as protein binders over traditional antibodies as they can be expressed intracellularly as “intrabodies” (Plückthun, 2015). Given their highly specific synaptic labelling, DARPin expression could be used as a tool to visualise inhibitory synapses in living neurons or non-neuronal cells *in vivo* after by fusing DARPin clones to genetically encoded fluorescent proteins. The small genetic size of DARPins allows for their packaging along with additional elements such as inducible expression systems or other functional moieties into viral vectors with small genomic packaging limits. Future derivatisation of anti-gephyrin DARPin binders, e.g. using cell type specific drivers to express DARPins fused to different genetically encoded fluorescent proteins, could improve our understanding of how the inhibitory postsynapse remodels similarly or differentially within excitatory and inhibitory neurons within the same circuit after experimental intervention.

### Use of anti-gephyrin DARPins as biochemical tools

While gephyrin is used experimentally to morphologically identify the inhibitory postsynapse, it achieves its function through protein-protein interactions. Unbiased protein interaction network identification broadens how we envisage protein function and regulation. For example a BioID-based gephyrin interactome discovered a novel inhibitory synaptic protein, InSyn1 (Uezu et al., 2016), which was found to be a key regulator of the dystroglycan complex and important for cognitive function (Uezu et al., 2019). By combining identified gephyrin interactors from antibody-based and DARPin-based experiments (including three distinct DARPin clones with different binding modalities), we were able to develop a consensus gephyrin interactome which facilitates higher confidence pursuit of understanding how these proteins integrate or are regulated by gephyrin function. The thresholds and criteria used to identify gephyrin interactors were established to be inclusive, and are indeed able to capture a majority of established canonical gephyrin interactors, yet further assessment will be required to determine which interactors are functional, and additionally whether they interact with synaptic versus non-synaptic gephyrin.

Various interactome determination techniques may capture different pictures of gephyrin protein networks. Proximity-ligation based methods require expression of recombinant bait protein, which may not correspond to the endogenous expression level or diversity of isoforms of native proteins in cells, though they are able to capture transient interactions (Burke et al., 2015). Affinity purification of gephyrin protein complexes is more likely to capture stable gephyrin protein complexes and may not identify transient interactors, but it allows for identification of native gephyrin protein complexes reflecting the heterogeneity in its isoforms present or its posttranslationally modified state. Therefore, using proximity-based labeling systems such as APEX, TurboID, et cetera in conjunction with DARPins will allow for a comparison of stable (possibly structural) functions of gephyrin and transient (possibly signaling) roles of gephyrin.

Within our interactome data, we found previously unidentified but presumed interacting proteins which are well known regulators of gephyrin. These include kinases such as CAMKIIα, which enhances gephyrin scaffolding via phosphorylation of serine 305 (Flores et al., 2015), GSK3β which phosphorylates serine 270, and MK01 (ERK2) which targets serine 268 to reduce clustering (Tyagarajan et al., 2013), as well as Protein Phosphatase 2A which antagonizes gephyrin phosphorylation at serine 270 (Kalbouneh et al., 2014). Additionally, we found the presence of multiple signaling scaffolds including CNKR2, a PSD-associated protein which may regulate RAS-dependent MAPK signaling and is associated with intellectual disability in humans (Hu et al., 2016). This protein was very recently confirmed to regulate network excitability using a genetic model (Erata et al., 2021). These data suggest that many of the kinases known to regulate gephyrin scaffolding as well as regulators of those kinases are part of gephyrin protein complexes. Discovering how these kinase scaffolds associate and regulate gephyrin via phosphorylation may pave the way for targeted therapeutic development.

The name “gephyrin” is derived from the Greek word γέφυρα meaning “bridge” as it was discovered to link glycine receptors to the cytoskeleton (P. Pfeiffer et al., 1982; Prior et al., 1992), and subsequently found to interact with other cytoskeletal components including dynein light chains 1 and 2 (Fuhrmann et al., 2002). We have now expanded this list to include multiple cytoskeletal interactors including those involved in microtubule nucleation during cell division (e.g. TBG1, CENPV). Interestingly, gephyrin colocalised with microtubule nucleation centres has been recently identified in U2OS cells (Zhou et al., 2021).

Our consensus interactome identified not only canonical gephyrin binders but also unexpected proteins related to mRNA regulation, metabolism, and ribosomal function which may suggest non-synaptic functions of gephyrin yet to be described, the signifiance of which can now be investiated further with independent methods. Canonical gephyrin interactors differed in their abundance within complexes precipitated by clones which bind the P1 or C3 cassette variants suggesting that different DARPin clones can access distinct synaptic gephyrin complexes. Gephyrin has been implicated previously in regulation of mTOR, a signaling scaffold (Machado et al., 2016; Sabatini et al., 1999; Wuchter et al., 2012) as well as with elongation factor EF1A1 which along with mTOR directs mRNA translation and acts as a cytoskeletal adaptor complex (Becker et al., 2013). We identified EF1A1 as an interactor enriched in DARPin-precipitated complexes along with other mRNA binding proteins involved in mRNA splicing and transport (e.g. PURA, PURB, PABP1). Additionally we detected the presence of transcription regulators such as SAFB1, DDX3X, and SIR2 from all DARPin complexes, and additional transcription factors including MECP2 (a Rett-syndrome associated protein regulating inhibitory network development (Pelkey et al., 2017) and present at the PSD (Aber et al., 2003)) found only in 27B3 and 27G2 gephyrin complexes. Gephyrin signaling has recently been implicated in coupling transcriptional signaling via ARX in pancreatic beta cells (Berishvili et al., 2017), and may therefore be involved in regulating additional transcriptional coupling in the brain via these described transcription factors. Many of the unexpected ribosomal and mRNA binding proteins were not detected in the control condition or using clone 27F3, suggesting that non-specific binding to these classes of proteins is not an intrinsic property of DARPins. Further studies using isoform-specific DARPin clones to capture gephyrin protein networks in neuronal compared to non-neuronal cells will clarify which protein interactors may be isoform or cell type specific. Indeed our group recently demonstrated gephyrin affects microglial reactivity and synapse stability after stroke (Cramer et al., 2022).

### Further applications of DARPins

Beyond morphological and biochemical applications, DARPin binders can be developed further as functional tools. To date no full-length experimentally-determined gephyrin structural information exists, possibly due to the instability of gephyrin’s C domain, making holo-gephyrin crystallisation difficult (Sander et al., 2013), and approaches to stabilize gephyrin for structure determination will be important to understand its structure-function relationship at the synapse (Fritschy et al., 2008). The stabilisation of target proteins for structure determination has been a major experimental application of DARPins (Batyuk et al., 2016; Tamaskovic et al., 2012; Wu et al., 2018). In this study we identified one DARPin clone (27F3) which binds only to the full length P1 isoform but not individual domains. Using structural biology to assess the interaction between DARPins and full-length gephyrin, we may not only be able to rationally engineer DARPins to achieve different binding functionality, but may also derive fundamental information about gephyrin’s form and function relationships, which would be essential for any future therapeutic efforts targeting gephyrin.

### Importance of protein binder development for neuroscience

Several synthetic protein binder scaffolds exist, including DARPins, nanobodies, anticalins, affibodies, and others (Harmansa & Affolter, 2018), providing a plethora of platforms to develop tools that detect or modify synaptic proteins, yet their application in neuroscience has lagged behind other fields. Of note, a fibronectin-based scaffold was used to generate intrabodies (termed FingRs by the authors) against gephyrin and the excitatory postsynaptic scaffold protein PSD-95 (Gross et al., 2013). This system has been used chiefly to label gephyrin clusters in living neurons (Crosby et al., 2019; Gross et al., 2016; Son et al., 2016; Uezu et al., 2016), but has been limited in its virus-based *in vivo* labeling and morphological detection of native gephyrin in tissue. Therefore, our DARPin-based toolset complements previously developed tools for live imaging, and future studies will test whether DARPins may be similarly used for native gephyrin tagging in living neurons.

Due to their stability and structure, DARPins are facile and inexpensive to produce and purify using simple bacterial systems and affinity resins. In addition, DARPins have relatively small sizes and defined sequences which makes them experimentally tractable. We have shown that developing multiple DARPins to examine gephyrin is a useful strategy for understanding the heterogeneity of its signaling and function, and similar strategies applied to other synaptic beyond gephyrin are likely to yield fruitful insights, as previously demonstrated with other systems (Plückthun, 2015). For synaptic biology, these DARPins offer an additional toolset which we hope will be expanded in the future so that excellent and well characterised binders are available to probe a multitude of targets with the goal of enhancing research efficiency and facilitating discoveries.

## Materials and methods

### Cloning and expression of gephyrin phosphorylation mutants

The principal (P1) rat isoform of gephyrin (referred to as wild-type, WT), or the P1 variant containing mutated serine to alanine (phospho-null) or serine to glutamic acid (phospho-mimetic) mutations at serines 268 and 270 have been described previously (Tyagarajan et al., 2013). Primers introducing a 5’ EcoRI restriction site upstream of a 2x GSSS linker sequence and 3’ KpnI site (*see primer table*) were used to amplify WT or mutated gephyrin before restriction digest and ligation into target vectors for recombinant bacterial expression and purification containing a 5’ His_8_ tag or His-Avi tag. *E. coli* BL21-DE3 Gold was transformed with the correct clones, and clones containing the His-Avi tag were transformed along with a plasmid encoding BirA for AviTag-specific biotin ligation. Bacteria were grown in THY media (20 g tryptone, 10 g yeast extract, 11 g HEPES, 5 g NaCl, 1 g MgSO_4_/L pH 7.4) containing ampicillin (100 µg/mL) and chloramphenicol (10 µg/mL) to ensure expression of both tagged Avi-gephyrin and BirA. Overnight 5 mL cultures were used to inoculate a 150 mL culture grown at 37°C and 250 rpm until an OD_600_ of 0.7 was reached. Induction and biotinylation was achieved by using a final concentration of 30 µM IPTG and 50 µM D-biotin (dissolved in 10 mM bicine buffer, pH 8.3). Protein induction proceeded for 6 hours before bacteria were pelleted.

Bacterial pellets were re-suspended in 15 mL lysis buffer (50 mM Trizma base, 120 mM NaCl, 0.5% NP-40) containing cOmplete Mini protease inhibitor cocktail (Roche) and DNAseI (Roche) before sonication on ice to release proteins. The lysate was pelleted at 20,000 *g* at 4°C for 15 minutes, and the cleared lysate was passed through 0.45 and 0.22 µm sterile filters. His_8_-tagged proteins were affinity purified on a 1 mL nickel agarose column (HIS-Select) using gravity flow. The lysate volume was passed 2x through the column then washed 1x with 6 column volumes of medium salt equilibration buffer (300 mM NaCl, 50 mM NaH_2_PO_4_, 10 mM imidazole, pH 8.0), then 1x with low-salt buffer (same with 100 mM NaCl), 1x with medium-salt buffer (300 mM NaCl), 1x with high-salt buffer (same with 500 mM NaCl), then 2x with medium-salt buffer (300 mM NaCl). Proteins were eluted in 4 mL elution buffer (equilibration buffer containing 250 mM imidazole) and dialysed in storage buffer (150 mM NaCl, 50 mM NaH_2_PO_4_, pH 7.5) using dialysis tubing. Dialysed protein was centrifuged at 60,000 *g* to remove any aggregated products, and the concentration was determined using absorption at 280 nm using a Nanodrop spectrophotometer with predicted protein molecular weight and extinction coefficient values determined using ProtParam online software (ProtParam, Swissprot, https://web.expasy.org/protparam/). Protein biotinylation was assessed using a streptavidin shift assay and stored at −80°C.

### Anti-gephyrin DARPin selection and screening

To generate DARPin binders, biotinylated gephyrin S268E/S270E was immobilized alternately on either MyOne T1 streptavidin-coated beads (Pierce) or Sera-Mag neutravidin-coated beads (GE), depending on the particular selection round. Ribosome display selections were performed essentially as described (Dreier & Plückthun, 2012), using a semi-automatic KingFisher Flex MTP96 well platform. The library includes N3C-DARPins with stabilized C-terminal caps (Kramer et al., 2010). This library is a mixture of DARPins with randomised and non-randomised N- and C-terminal caps respectively (Plückthun, 2015; Schilling et al., 2014), and successively enriched pools were cloned as intermediates in a ribosome display specific vector (Schilling et al., 2014). Selections were performed over four rounds with decreasing target concentration and increasing washing steps to enrich for binders with high affinities. The first round included the initial selection against gephyrin S268E/S270E at low stringency. The second round included pre-panning with the opposite phospho-null (gephyrin S268A/S270A) variant immobilized on magnetic beads, with the supernatant transferred to immobilized target of the same variant. The 3^rd^ round included this pre-panning of the opposite variant and the addition of the (non-biotinylated) same variant to enrich for binders with slow off rate kinetics. The 4^th^ and final round included only the pre-panning step and selection was performed with low stringency.

The final enriched pool was cloned as fusion construct into a bacterial pQE30 derivative vector with a N-terminal MRGS(H)_8_ tag (His_8_) and C-terminal FLAG tag via unique BamHI x HindIII sites containing *lacIq* for expression control. After transformation of *E. coli* XL1-blue, 380 single DARPin clones for each target protein were expressed in 96 well format and lysed by addition of a (concentrated Tris-HCL based HT-Lysis buffer containing octylthioglucoside (OTG), lysozyme and nuclease or B-Per Direct detergent plus lysozyme and nuclease, Pierce). These bacterial crude extracts of single DARPin clones were subsequently used in a Homogeneous Time Resolved Fluorescence (HTRF)-based screen to identify potential binders. Binding of the FLAG-tagged DARPins to streptavidin-immobilized biotinylated Gephyrin variants was measured using FRET (donor: streptavidin-Tb, acceptor: anti-FLAG-d2, Cisbio). Further HTRF measurement against ‘No Target’ allowed for discrimination of Gephyrin-specific hits.

From the identified binders, 32 were sequenced and 25 unique clones were identified. The DARPins were expressed in small scale, lysed with Cell-Lytic B(SIGMA) and purified using a 96 well IMAC column (HisPur^TM^ Cobalt plates, Thermo Scientific). DARPins after IMAC purification were analyzed at a concentration of 10 µM on a Superdex 75 5/150 GL column (GE Healthcare) using an Aekta Micro system (GE Healthcare) with PBS containing 400 nM NaCl as the running buffer to identify monomeric DARPin binders. Final hit validation of specificity was performed by ELISA using small scale IMAC-purified DARPins. Binding of the FLAG-tagged DARPins to streptavidin-immobilized biotinylated gephyrin variants was measured using a mouse-anti-FLAG-M2 antibody (Sigma) as 1^st^ and goat-anti-mouse-alkaline phosphatase conjugated antibody (Sigma) as 2^nd^ antibody. Further ELISA measurement against ‘No Target’ allowed for discrimination of Gephyrin-specific hits. The best binders did not descrimination between phospho-mimetic states, suggesting that other epitopes were favoured.

### Cloning and recombinant expression of anti-gephyrin DARPins

Bacterial expression and purification of FLAG-tagged DARPins was performed as for His-tagged gephyrin constructs. Purification was validated using SDS-PAGE and Coomassie staining of acrylamide gels. Sub-cloning of select DARPins into a vector containing an N-terminal HSA leader sequence and C-terminal human Fc fragment (hFc) region using BamHI and HindIII restriction sites was performed for mammalian cell production. Test rounds of DARPin-hFc fusion expression were performed in adherent HEK293T cells where the supernatant was collected to confirm DARPin hFc expression. Medium-scale production of DARPin-hFc fusion constructs was performed with assistance from the Protein Production and Structure core facility (PTPSP Lausanne) by transfecting plasmids for clones 27B3-hFc, 27F3-hFc, and 27G2-hFc as well as control DARPin E3_5-hFc into non-adherent HEK cells and grown in 400 mL cultures. DARPin-hFc recombinant protein was affinity-purified using Protein A resin after overnight incubation with rotation at 4°C, and captured on a 15 mL column Protein A Sepharose resin (Genscript), beads were washed with 50 column volumes of PBS and eluted with glycine buffer pH 3.0 into 1.5 M Tris-HCl pH 8.0 before overnight dialysis into PBS pH 7.5. Concentration was determined using a Nanodrop spectrophotometer using the A280 extinction coefficient.

### Gephyin binding fluorescence assay in HEK293T cells

An in-cell fluorescence-based assay was developed to characterize the relative binding of anti-gephyrin DARPin clones to eGFP-tagged gephyrin variants in order to assess binding and to validate the DARPin screening ELISA results in cells. HEK293T cells were maintained in DMEM with 10% FCS at 37°C in a 5% CO_2_ jacketed incubator. Cells were seeded onto glass coverslips and grown to 50% confluency before transfecting plasmids (using standard PEI-based transfection at a ratio of 1 µg plasmid to 4 µg PEI). eGFP-tagged gephyrin P1 variant, as well as those containing serine-to-alanine or -glutamate mutations at S268 and S270 (S268A/S270A, S268E/S270E) have been previously described (Tyagarajan et al., 2013). eGFP-tagged gephyrin E domain or GC domains (Lardi-Studler et al., 2007) as well as variants containing the C3 or C4a splice cassettes (Lardi-Studler et al., 2007) have been described previously. Cells grown on coverslips were washed briefly in PBS and fixed in 4% PFA for 15 minutes. Coverslips were washed in PBS, then treated with 1:2000 (1mg/ml stock) dilution of DARPin-FLAG clones or a control clone (non-binding DARPin E3_5-FLAG) in 10% normal goat serum (NGS) for 90 minutes. Coverslips were washed and then treated with a 1:1000 dilution of mouse anti-FLAG antibody (clone M2, Sigma) for 60 minutes then washed 3x in PBS. Coverslips were incubated with an Alexa 647-conjugated goat anti-mouse secondary antibody and DAPI for 30 minutes prior to washing 3x with PBS and drying before mounting with DAKO mounting medium onto glass slides.

Coverslips were imaged using an LSM700 microscope (Zeiss) with 40x (1.4 NA) objectives. Images were acquired using Zen software (Zeiss). Laser intensity and gain settings were set to maximize signals in all channels/conditions without bleed-through or signal saturation, and acquisition settings were kept consistent for comparative analyses. eGFP-gephyrin-positive HEK cells were imaged at random locations on the coverslip, and fluorescent signals were acquired at 8 bits in the 488 and 647 channels to capture the eGFP-gephyrin and FLAG signal, respectively. eGFP-gephyrin presents as a diffuse signal in the soma with occasional cytoplasmic aggregates. For intensity analysis, ROIs were manually drawn within the cytosol to avoid inclusion of these aggregates in the quantification. Fluorescence intensity was quantified using ImageJ. The slope of the relationship between the eGFP-gephyrin signal and the FLAG signal was used to compare relative binding of DARPins to their target.

### Animals

All procedures fulfilled the ARRIVE guidelines on experimental design, animal allocation to different experimental groups, blinding of samples to data analysis and reporting of animal experiments. We conducted a sample size calculation based on previous experiments for synaptic analysis with effect size of 0.2, a power of 0.8, and a significance level of 0.05. The data in our study included 5-6 animals per genotype, which exceeded the sample size calculation. Randomization of experimental cohorts is achieved by separating the age matched animals into male and female sexes to ensure that both genders are equally represented in the experimental groups. The experimenter is blinded to the experiments by another student assigning numbers and allocating animals to different groups at the start. https://www.isogenic.info/html/7__randomisation.html#methods

C56Bl/6J mice were purchased from Charles River (Germany) and timed-pregnant Wistar rats (for E17 embryo collection for neuron culture) were purchased from Envigo (Netherlands). The S268A/S270A phospho-null mouse was previously generated using CRISPR-Cas9 editing to mutate residues at the endogenous locus (Cramer et al., 2022). The collection of embryonic and adult tissue was performed in accordance with the European Community Council Directives of November 24^th^ 1986 (86/609/EEC). Tissue collection was performed under license ZH011/19 approved by the Cantonal Veterinary office of Zurich.

### Synaptic staining, imaging, and analysis

Hippocampal cell cultures derived from E17 Wistar rat embryos were prepared as previously described (Tyagarajan et al., 2013) containing a mixture of excitatory/inhibitory neurons and glia grown on poly-L-lysine-coated glass coverslips. Cultures were maintained for 15 days in vitro (DIV) before use to allow for synapse formation. Neurons were prepared for DARPin-FLAG or DARPin-hFc staining and immunostaining as with HEK293T cultures, with the exception that endogenous gephyrin was analysed using the anti-gephyrin antibody clone Ab7a (Sysy 147 011) or clone 3B11 (Sysy 147 111). Guinea pig anti-VGAT antibody (Sysy 131 004) and mouse anti-Ankyrin G (Neuromab, MABN466) were used to identify inhibitory presynapses and the axon initial segment, respectively. Homemade affinity purified guinea pig anti-GABRA2 was used to detect post-synaptic sites in tissue. Optimal concentrations of anti-gephyrin DARPins for staining were determined for each clone, 1:2000 dilution from 1 mg/mL stock was determined to be best for DARPin-FLAG, 1:4000 dilution performed best for DARPin-hFc.

For brain tissue staining, animals were anesthetised with intraperitoneal injections of pentobarbital before trans-cardial perfusion with oxygenated, ice cold artificial cerebrospinal fluid (ACSF: 125 mM NaCl, 2.5 mM KCl, 1.25 mM NaH_2_PO_4_, 26 mM NaHCO_3_, 25 mM D-glucose, 2.5 mM CaCl_2_, and 2 mM MgCl_2_). Perfused brains were dissected and post-fixed in 150 mM phosphate-buffered saline (PBS) containing 4% paraformaldehyde (PFA) (pH 7.4) for 90 minutes at 4 °C. Tissue was cryoprotected overnight in PBS containing 30% sucrose 4 °C, then cut into 40 µm thick sections using a sliding microtome. Sections were stored at −20 °C in antifreeze solution (50 mM sodium phosphate buffer with 15% glucose, 30% ethylene glycol at pH 7.4) until use. For immunofluorescence experiments, sections were washed 3 x 10 minutes under gentle agitation in TBST (50 mM Tris, 150 mM NaCl, 1% Tween, pH 7.5) before overnight incubation in primary antibody solution (with or without DARPin inclusion) (TBST containing 0.2% Triton X-100 and 2% NGS). For DARPin-hFc 27G2, a concentration of 1:4000 was used (from 1 mg/mL stock). Sections were then washed 3 x 10 minutes and incubated for 30 minutes at room temperature with secondary antibodies in TBST solution with 2% normal goat serum NGS (Jackson). Sections were washed again 3 x 10 minutes in TBST before transfer to PBS and mounting onto gelatine-coated slides, then covered using DAKO mounting medium. For all tissue morphological analysis, image acquisition, processing, and analysis was acquired/performed blind to condition using identical imaging parameters. Images used for synapse quantification experiments were acquired on a Zeiss LSM 800 laser scanning confocal microscope operating Zen image acquisition software (Zen 2011) using 63x oil immersion objectives (N.A. 1.4). Identical imaging settings were used when comparing between groups in a given experiment. Relative Ab7a/DARPin-hFc 27G2 fluorescent intensity cluster analysis was performed using the Analyse Particles functionality of FIJI after thresholding. Synaptic colocalisation analysis was performed using a custom ImageJ macro previously described (Panzanelli et al., 2017).

### Precipitation of gephyrin complexes for LC-MS/MS interactome determination

Tissue lysates were prepared from acutely isolated cortexes and hippocampi of 4 male and 4 female C57BL/6J mice (Charles River) on ice and immediately homogenized in cold EBC lysis buffer (50 mM Tris-HCl, 120 mM NaCl, 0.5% NP-40, and 5 mM EDTA with cOmplete mini protease inhibitors (Roche) and phosphatase inhibitor cocktails 2 and 3 (Sigma)) and incubated on ice for 60 minutes. Lysates were cleared by centrifugation at 20,000 *g* for 20 minutes and the supernatant protein concentration measured using a BCA assay. Gephyrin complexes were captured by incubating protein lysate (total 6 mg of protein per reaction) with DARPin-hFc binders or the control DARPin clone E3_5 or, control IgG, or 3B11 mouse-anti-gephyrin antibody for 3 hours at 4° C with rotation. In order to precipitate similar amounts of gephyrin protein, 4 µg of 3B11 antibody, or approximately 2 µg of anti-gephyrin DARPin-hFc (adjusted for equimolar concentration) were used per reaction (1.5 mL volume total). Complexes were precipitated using 20 µg of Protein G magnetic beads (30 minutes incubation with rotation), and washed 6x in 600 µl of EBC buffer. The supernatant was removed and replaced with 25 µl of PBS and immediately submitted for LC-MS/MS sample preparation.

### Immunoblotting

For immunoblotting experiments, input and precipitated samples were prepared in 5x SDS buffer containing beta-mercaptoethanol (Bio-Rad) and boiled for 5 minutes at 90° C. Protein concentration determination was performed using a BCA assay (Pierce). Acrylamide gels were either stained with Coomassie dye or transferred to PVDF membranes. Gephyrin was detected using a mouse anti-gephyrin antibody (clone 3B11, 1:1,000), and DARPin-hFc was detected using an anti-hFc (HRP conjugated, 1:40,000) antibody overnight and detected using anti-mouse IR 680 dye (LI-COR) on a LI-COR imager, or an HRP detection kit using a Fuji imager.

### On bead digestion

Captured immunocomplexes were processed immediately after precipitation. Beads were washed once in 100 µL digestion buffer (10 mM Tris + 2 mM CaCl_2_, pH 8.2). After resuspension in 45 µl digestion buffer, proteins were reduced and alkylated with 2 mM TCEP and 20 mM chloroacetamide, respectively, for 30 min at 60 °C in the dark. Five µL of Sequencing Grade Trypsin (100 ng/µl in 10 mM HCl, Promega) were added to the beads and the digestion was carried out in a microwave instrument (Discover System, CEM) for 30 min at 5 W and 60 °C. The supernatants were transferred into new tubes and the beads were washed with 150 µl 0.1% TFA then pooled with the previous supernatant. The samples were dried and re-solubilized with 20 µl of 3% acetonitrile, 0.1% formic acid for MS analysis. Prior to MS analysis, the peptides were diluted to an absorption (A280) of 0.2.

### Liquid chromatography-mass spectrometry analysis

Mass spectrometry analysis was performed on an Orbitrap Fusion Lumos (Thermo Scientific) equipped with a Digital PicoView source (New Objective) and coupled to a M-Class UPLC (Waters). Solvent composition at the two channels was 0.1% formic acid for channel A and 0.1% formic acid, 99.9% acetonitrile for channel B. For each sample 1 μL of diluted peptides were loaded on a commercial MZ Symmetry C18 Trap Column (100 Å, 5 µm, 180 µm x 20 mm, Waters) followed by nanoEase MZ C18 HSS T3 Column (100 Å, 1.8 µm, 75 µm x 250 mm, Waters). The peptides were eluted at a flow rate of 300 nL/min using a gradient from 5 to 22% B in 80 min, 32% B in 10 min and 95% B for 10 min. The mass spectrometer was operated in data-dependent mode (DDA) acquiring a full-scan MS spectra (300−1,500 m/z) at a resolution of 120,000 at 200 m/z after accumulation to a target value of 500,000. Data-dependent MS/MS spectra were recorded in the linear ion trap using quadrupole isolation with a window of 0.8 Da and HCD fragmentation with 35% fragmentation energy. The ion trap was operated in rapid scan mode with a target value of 10,000 and a maximum injection time of 50 ms. Only precursors with intensity above 5,000 were selected for MS/MS and the maximum cycle time was set to 3 s. Charge state screening was enabled. Singly, unassigned, and charge states higher than seven were rejected. Precursor masses previously selected for MS/MS measurement were excluded from further selection for 20 s, and the exclusion mass tolerance was set to 10 ppm. The samples were acquired using internal lock mass calibration on m/z 371.1012 and 445.1200. The mass spectrometry proteomics data were handled using the local laboratory information management system (LIMS) (Türker et al., 2010).

### Protein identification and label-free protein quantification

The acquired raw MS data were processed by MaxQuant (version 2.0.1.0), followed by protein identification using the integrated Andromeda search engine (Cox & Mann, 2008). Spectra were searched against a Uniprot *Mus musculus* reference proteome (taxonomy 10090, version from 2019-07-09), concatenated to its reversed decoyed FASTA database and common protein contaminants. Carbamidomethylation of cysteine was set as fixed modification, while methionine oxidation, STY phosphorylation and N-terminal protein acetylation were set as variable. Enzyme specificity was set to trypsin/P allowing a minimal peptide length of 7 amino acids and a maximum of two missed cleavages. The maximum false discovery rate (FDR) was set to 0.01 for peptides and 0.05 for proteins. Label-free quantification was enabled and a 2-minute window for match between runs was applied. In the MaxQuant experimental design template, each file is kept separate in the experimental design to obtain individual quantitative values. Protein fold changes were computed based on Intensity values reported in the proteinGroups.txt file. A set of functions implemented in the R package SRMService (W. Wolski, J. Grossmann, C. Panse. 2018. SRMService - R-Package to Report Quantitative Mass Spectrometry Data. http://github.com/protViz/SRMService) was used to filter for proteins with 2 or more peptides allowing for a maximum of 3 missing values, and to compute p-values using the t-test with pooled variance. If all measurements of a protein are missing in one of the conditions, a pseudo fold change was computed, replacing the missing group average by the mean of the 10% smallest protein intensities in that condition. To determine DARPin and GEPH isoform coverage in the individual pulldown conditions, the data were processed and searched with Proteome Discoverer 2.5 using Sequest and Percolator with Protein Grouping deactivated and only unique peptides were used for quantification.

### Interactome analysis

Proteins were considered present when detected using at least 2 unique peptide signatures in all replicates of a given binder. Interactors were considered part of gephyrin complexes when either 1) not present in the control condition, or 2) enriched by a log2 fold-change in abundance of at least 2.5 in the binder condition with an FDR cut-off of 0.05. These thresholds allowed for complete coverage of known gephyrin interactors. Binders common to multiple interactomes were identified using Microsoft Excel for comparison of ontology and abundances. Venn diagrams were visualized using InteractiVenn (http://www.interactivenn.net/). Protein ontology was identified and grouped, and enrichment determined using WebGestalt over-representation analysis (http://www.webgestalt.org/), Gene Ontology Resource identification (http://geneontology.org/), and Uniprot (https://www.uniprot.org/). Interaction networks were generated using STRING version 11.5 and imported to Cystoscape version 3.8.2 for visualisation. Network map edges represent putative relationships between protein nodes as identified by STRING. Node size is colored based on functional ontology, and size based on abundance relative to gephyrin in each experiment. Canonical gephyrin interactors include Collybistin (ARGH9), GABA_A_R subunits (GBRA1, 2,3, GABG2, GBRB2, 3), glycine receptor subunits (GLRB, GLRA), dynein light chain (DYL1, 2), IQSEC3 (IQEC3), Dystrobrevin alpha (DNTA), Ena VASP-like (EVL), MENA (ENAH), the proline cis-trans isomerase PIN1, profilins 1 and 2 (PROF1, 2), neuroligin 2 (NLGN2), reviewed in (Groeneweg et al., 2018). Protein names used for display are the official Uniprot protein ID designation. Uniprot protein IDs were used for cross-experiment comparison and ontology searches.

### Statistical tests

Statistical tests and significance are reported in the figure captions. Statistical analysis was performed using Microsoft Excel and Graph Pad Prism 8.0. Normality tests were performed on data to evaluate correct application of parametric or non-parametric analysis, with the exception of experiments using small sample sizes (n<4) where parametric comparisons were used.

### Visual representation

Data plots were generated using Microsoft Excel or GraphPad Prism 8. Images were visualized and processed in FIJI (1.53q). Images brightness was enhanced for display by adjusting the brightness and contrast for display purposes, but when comparing between experimental conditions, all images were enhanced with the same settings to preserve apparent differences in morphology and intensity. Diagrams and figures were arranged in InkScape (version 1.0), and text and tables were arranged using the Microsoft Office Suite. Sequence alignment was performed using ClustalW and visualized using JalView. Heat map generation and hiearchical clustering was performed with Morpheus (https://software.broadinstitute.org/morpheus).

## Material availability

The use of the anti-gephyrin DARPin constructs presented in this manuscript will be made available following an academic use MTA agreement.

## Data availability

All relevant mass spectrometry data has been deposited to the ProteomeXchange Consortium via the PRIDE (http://www.ebi.ac.uk/pride) partner repository.

**Project Name:** Gephyrin interactome from mouse brain lysates using anti-gephyrin antibody and anti-gephyrin DARPins

**Project accession:** PXD033641

**Project DOI:** 10.6019/PXD033641

## Key resources tables

**Table 1.** List of plasmids used in this study. RRIDs given where available. NA: not applicable.

| Reagent type (species or resource) | Designation | Source or reference | Identifiers | Additional information |
| --- | --- | --- | --- | --- |
| Plasmid backbone | GST within 3' 6x His Tag | Provided by the UZH High Throughput Binder Selection platform | pET20b-A(H6)-GST | Used for subcloning recombinant gephyrin constructs for recombinant bacterial expression for use in the ribosome display selection. |
| Plasmid backbone | GST within 3' 6x His Tag and Avi tag | Provided by the UZH High Throughput Binder Selection platform | pET20b-A(H6)-AviTag | Used for subcloning recombinant gephyrin constructs for recombinant bacterial expression for use in the ribosome display selection. |
| Plasmid | BirA enzyme | Provided by the UZH High Throughput Binder Selection platform | pBirAcm | Encodes the AVI-tag specific biotin ligase BirA for biotin-tagging of recombinant gephyrin constructs for use in the ribosome display selection. |
| Plasmid backbone | N-terminal 8xHis tag and C-terminal FLAG tag bacterial expression vector | Provided by the UZH High Throughput Binder Selection platform | pQIq_MRGS_HIS8_(DARPin)_FLAG | Used as the backbone for inserting DARPins using HindIII and BamHI restriction sites for recombinant bacterial expression of FLAG tagged DARPins. |
| Plasmid backbone | N-terminal HSA leader sequence and C-terminal hFc tag for mammalian expression | Provided by the UZH High Throughput Binder Selection platform | pcDNA3.1_SacB_hFc | Used as the backbone for inserting DARPins using HindIII and BamHI restriction sites for recombinant mammalian expression of hFc tagged DARPins. |
| Plasmid | N-terminal His-tagged P1-gephyrin S268/270A | This article | pET20b-A(H6)- P1-gephyrin S268/270A | Subcloned from pEGFPC2-gephyrin S268/270A (Tyagarajan et al., 2013) using added Kpn1 and EcoRI sites into pET20b-A(H6)-GST for use in DARPin ribosome display selection. |
| Plasmid | N-terminal His-tagged P1-gephyrin S268/270E | This article | pET20b-A(H6)- P1-gephyrin S268/270E | Subcloned from pEGFPC2-gephyrin S268/270E (Tyagarajan et al., 2013) using added Kpn1 and EcoRI sites into pET20b-A(H6)-GST for use in DARPin ribosome display selection. |
| Plasmid | N-terminal His-tagged P1-gephyrin | This article | pET20b-A(H6)- P1-gephyrin | Subcloned from pEGFPC2-gephyrin P1 (Tyagarajan et al., 2013) using added Kpn1 and EcoRI sites into pET20b-A(H6)-GST for use in DARPin ribosome display selection. |
| Plasmid | N-terminal HisAvi-tagged P1-gephyrin S268/270A | This article | pET20b-A(H6)- P1-gephyrin S268/270A AviTag | Subcloned from pEGFPC2-gephyrin S268/270A (Tyagarajan et al., 2013) using added Kpn1 and EcoRI sites into pET20b-A(H6)-AviTag for use in DARPin ribosome display selection. |
| Plasmid | N-terminal HisAvi-tagged P1-gephyrin S268/270E | This article | pET20b-A(H6)- P1-gephyrin S268/270E AviTag | Subcloned from pEGFPC2-gephyrin S268/270E (Tyagarajan et al., 2013) using added Kpn1 and EcoRI sites into pET20b-A(H6)-AviTag for use in DARPin ribosome display selection. |
| Plasmid | N-terminal eGFP-tagged P1-gephyrin S268/270A | (Tyagarajan et al., 2013) | pEGFPC2-gephyrin S268/270A | Used for subcloning for recombinant bacterial expression as well as the in-cell fluorescence assays. |
| Plasmid | N-terminal eGFP-tagged P1-gephyrin S268/270E | (Tyagarajan et al., 2013) | pEGFPC2-gephyrin S268/270E | Used for subcloning for recombinant bacterial expression as well as the in-cell fluorescence assays. |
| Plasmid | N-terminal eGFP-tagged P1-gephyrin | (Tyagarajan et al., 2013) | pEGFPC2-gephyrin P1 | Used for subcloning for recombinant bacterial expression as well as the in-cell fluorescence assays. |
| Plasmid | N-terminal eGFP-tagged gephyrin GC domain | (Lardi-Studler et al., 2007) | EGFPC2-Gephyrin GC | Used for in cell fluorescence assays to assess relative binding of DARPins to the GC domain of gephyrin. |
| Plasmid | N-terminal eGFP-tagged gephyrin E domain | (Lardi-Studler et al., 2007) | EGFPC2-Gephyrin E | Used for in cell fluorescence assays to assess relative binding of DARPins to the E domain of gephyrin. |
| Plasmid | N-terminal eGFP-tagged gephyrin containing the C3 cassette | (Smolinsky et al., 2008) | pEGFPC2 Gephyrin C3 | Used for in cell fluorescence assays to assess relative binding of DARPins to the C3 cassette containing gephyrin variants. |
| Plasmid | N-terminal eGFP-tagged gephyrin containing the C4a cassette | (Smolinsky et al., 2008) | pEGFPC2 Gephyrin C4a | Used for in cell fluorescence assays to assess relative binding of DARPins to the C4a cassette containing gephyrin variants. |
| Plasmid | DARPin-FLAG E3_5 (control) | This article | pQlq_MRGS_HIS8_(E3_5)_FLAG | Created by subcloning DARPin E3_5 into pQlq_MRGS_HIS8_(DARPin)_FLAG using BamHI and HindIII sites. |
| Plasmid | DARPin-FLAG 27B3 | This article | pQlq_MRGS_HIS8_(27B3)_FLAG | Created by subcloning DARPin 27B3 into pQlq_MRGS_HIS8_(DARPin)_FLAG using BamHI and HindIII sites. |
| Plasmid | DARPin-FLAG 27D3 | This article | pQlq_MRGS_HIS8_(27D3)_FLAG | Created by subcloning DARPin 27D3 into pQlq_MRGS_HIS8_(DARPin)_FLAG using BamHI and HindIII sites. |
| Plasmid | DARPin-FLAG 27F3 | This article | pQlq_MRGS_HIS8_(27F3)_FLAG | Created by subcloning DARPin 27F3 into pQlq_MRGS_HIS8_(DARPin)_FLAG using BamHI and HindIII sites. |
| Plasmid | DARPin-FLAG 27B5 | This article | pQlq_MRGS_HIS8_(27B5)_FLAG | Created by subcloning DARPin 27B5 into pQlq_MRGS_HIS8_(DARPin)_FLAG using BamHI and HindIII sites. |
| Plasmid | DARPin-FLAG 27D5 | This article | pQlq_MRGS_HIS8_(27D5)_FLAG | Created by subcloning DARPin 27D5 into pQlq_MRGS_HIS8_(DARPin)_FLAG using BamHI and HindIII sites. |
| Plasmid | DARPin-FLAG 27G2 | This article | pQlq_MRGS_HIS8_(27G2)_FLAG | Created by subcloning DARPin 27G2 into pQlq_MRGS_HIS8_(DARPin)_FLAG using BamHI and HindIII sites. |
| Plasmid | DARPin-FLAG 27H2 | This article | pQlq_MRGS_HIS8_(27H2)_FLAG | Created by subcloning DARPin 27H2 into pQlq_MRGS_HIS8_(DARPin)_FLAG using BamHI and HindIII sites. |
| Plasmid | DARPin-FLAG 27G4 | This article | pQlq_MRGS_HIS8_(27G4)_FLAG | Created by subcloning DARPin 27G4 into pQlq_MRGS_HIS8_(DARPin)_FLAG using BamHI and HindIII sites. |
| Plasmid | DARPin-hFc E3_5 (control) | This article | pcDNA3.1_E3_5_hFc | Created by subcloning DARPin E3_5 into pcDNA3.1_SacB_hFc using BamHI and HindIII sites. |
| Plasmid | DARPin-hFc 27B3 | This article | pcDNA3.1_27B3_hFc | Created by subcloning DARPin 27B3 into pcDNA3.1_SacB_hFc using BamHI and HindIII sites. |
| Plasmid | DARPin-hFc 27F3 | This article | pcDNA3.1_27F3_hFc | Created by subcloning DARPin 27F3 into pcDNA3.1_SacB_hFc using BamHI and HindIII sites. |
| Plasmid | DARPin-hFc 27G2 | This article | pcDNA3.1_27G2_hFc | Created by subcloning DARPin 27G2 into pcDNA3.1_SacB_hFc using BamHI and HindIII sites. |

**Table 2.**
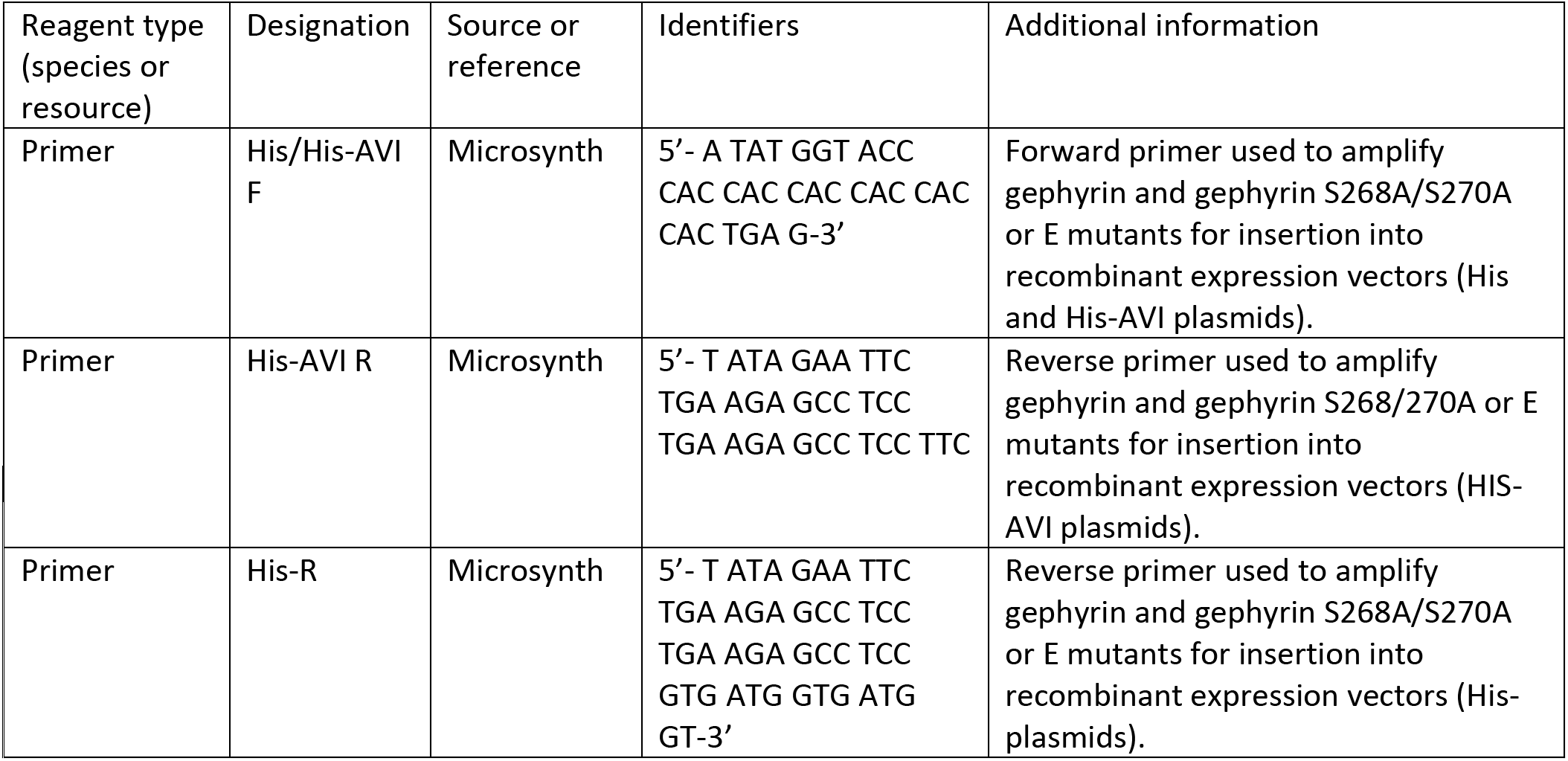
List of primers.

**Table 3.** List of antibodies/ protein binders and concentrations used. Dilution values correspond to manufacturer recommended reconstitution concentrations. Unless otherwise stated, stocks are 1 mg/ mL. RRIDs given where available. NA: not applicable.

| Reagent type (species or resource) | Designation | Source or reference | Identifiers | Additional information |
| --- | --- | --- | --- | --- |
| Primary antibody | Mouse anti-Ankyrin G (AnkG) | Neuromab | MABN466, RRID AB_274980 | IF/ICC used at 1:1000 |
| Primary antibody | Goat anti-mouse AP | Sigma-Aldrich (Merck) | A3562, AB_258091 | Used for ELISA screen |
| Primary antibody | Mouse anti-FLAG M2 | Sigma-Aldrich (Merck) | F3165, RRID AB_259529 | IF/ICC used at 1:1000 |
| Primary antibody | Mouse anti-FLAG D2 | Cisbio | 61FG2DLB | Used for HTRF screen. |
| Primary antibody | Guinea pig anti-GABRA2 | In house (J. -M Fritschy & Möhler, 1995) | - | IF/ICC used at 1:2000 |
| Primary antibody | Mouse anti-gephyrin 3B11 | Synaptic Systems | Cat #: 147111, RRID: AB_887719 | IF/ICC used at 1:1000 |
| Primary antibody | Rabbit anti-gephyrin Ab7a | Synaptic Systems | 147 008, RRID AB_2619834 | IF/ICC used at 1:2000 |
| Primary antibody | Guinea pig anti-VGAT | Synaptic Systems | 131308, AB_2832243 | IF/ICC used at 1:2000 |
| Secondary antibody | Goat anti-mouse Alexa Cy3 | Jackson ImmunoResearch Labs | JAC 115-165-166, RRID AB_2338692 | IF/ICC used at 1:500 |
| Secondary antibody | Goat anti-rabbit Alexa 488 | Jackson ImmunoResearch Labs | JAC 111-545-144, RRID AB_2338052 | IF/ICC used at 1:500 |
| Secondary antibody | Goat anti-Guinea pig Alexa 647 | Jackson ImmunoResearch Labs | JAC 106-605-003, RRID AB_2337446 | IF/ICC used at 1:500 |
| Secondary antibody | Goat anti-human Cy3 | Jackson ImmunoResearch Labs | JAC 109-165-170, AB_2810895 | IF/ICC used at 1:500 |
| Streptavidin conjugate | Streptavidin-Tb cryptate | Cisbio | 610SATLB | Used for HTRF screen. |
| Secondary antibody | IRDye 680RD Donkey anti-Mouse IgG | LI-COR Biosciences | LIC925-68072 | WB 1:20000 |
| HRP conjugate | Anti-human Fc HRP | CalBiochem | 401455 | WB 1:40000 |

**Table 4.** List of animal strains. RRIDs given where available. NA: not applicable.

| Reagent type (species or resource) | Designation | Source or reference | Identifiers | Additional information |
| --- | --- | --- | --- | --- |
| Rattus norvegicus | Wister rat (RccHan:WIST) | Envigo (Netherlands) | Order code: 168 | E17 embryos were collected from time mated dams. |
| Mus musculus | C57BL/6JCrI | Charles River Laboratories (Germany) | RRID IMSR_JAX:000664, | Used for synapse analysis and proteomic analysis. |
| Mus musculus | C57Bl6/JCrI GphnS268A/S270A | (Cramer et al., 2022) | NA | Used for synapse analysis only. |

**Table 5.** List of cell lines used in this study. RRIDs given where available. NA: not applicable.

| Reagent type (species or resource) | Designation | Source or reference | Identifiers | Additional information |
| --- | --- | --- | --- | --- |
| Cell line | BL21 DE3 Gold | BioRad | Cat #: 161-0156 | Used for recombinant bacterial gephyrin and DARPin expression. |
| Cell line | E.coli XL1-blue | Agilent | 200249 | Used for DARPin ribosome display screening. |
| Cell line | HEK293T | ATCC | CRL 11268 | Used for in cell DARPin binding screen. |

**Table 6.**
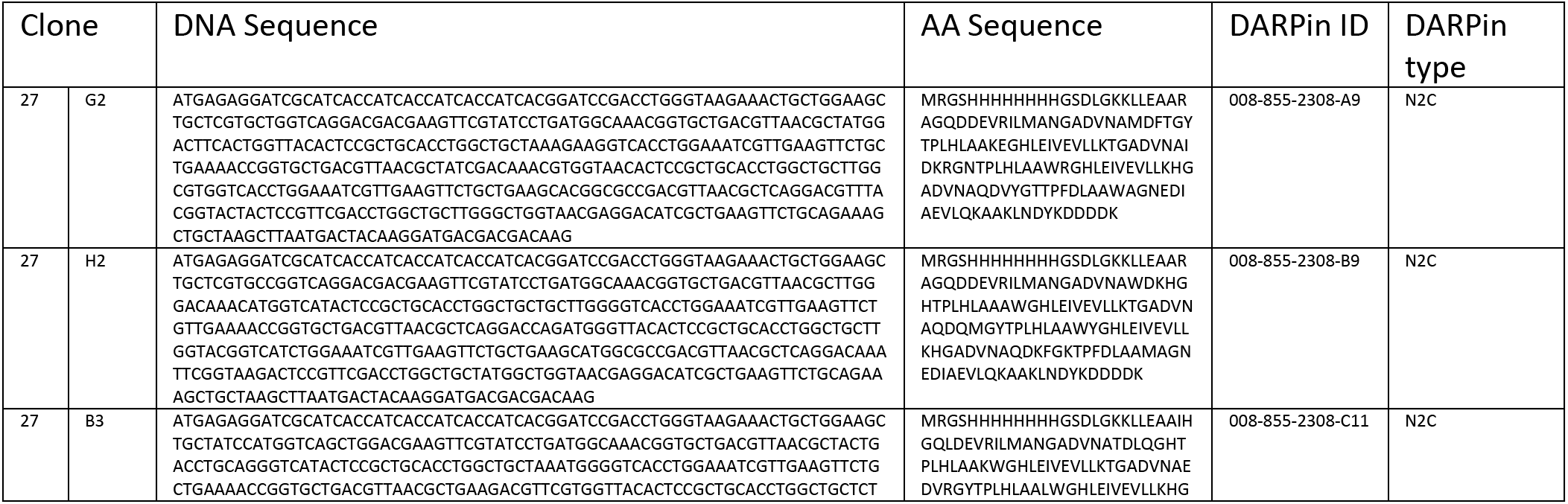
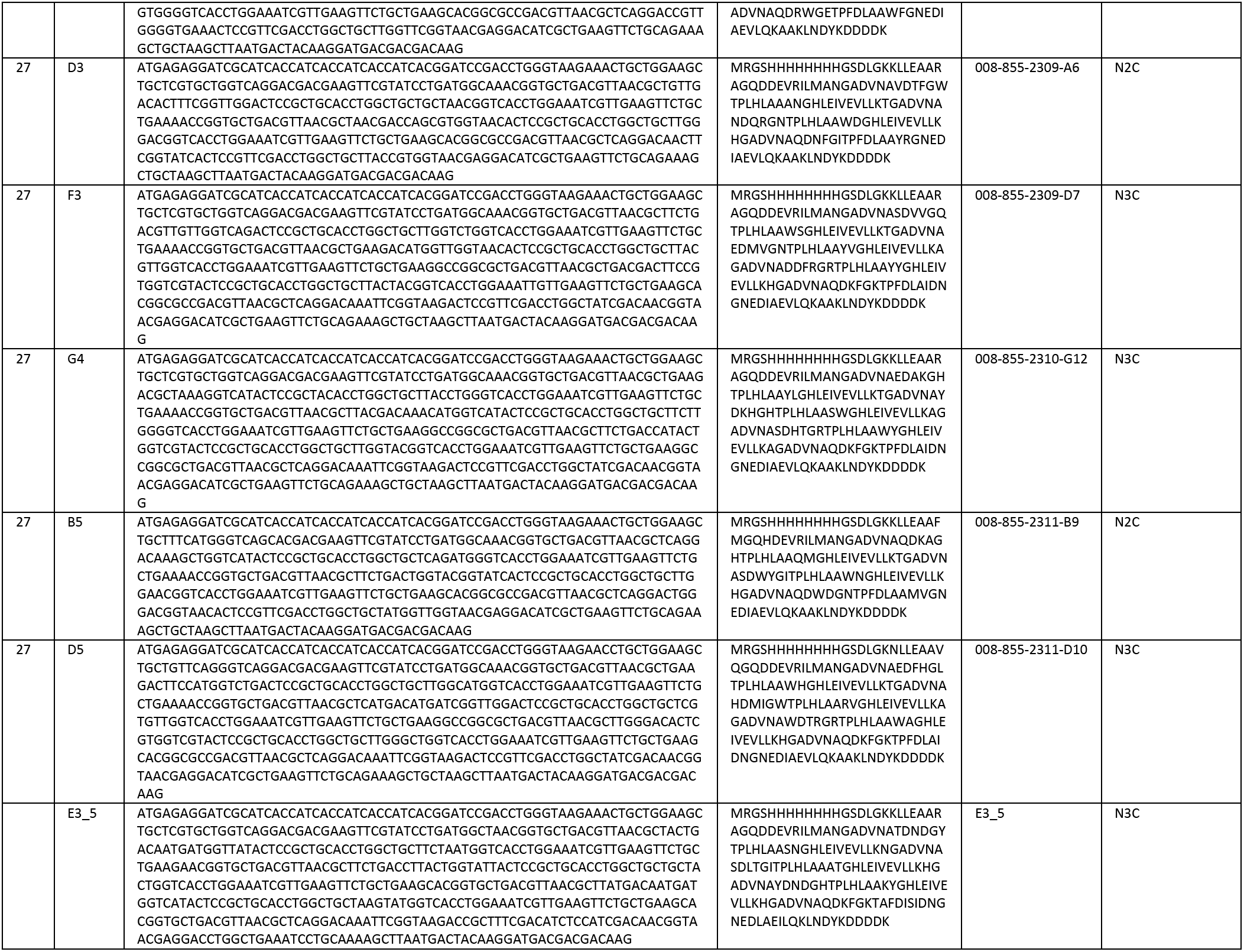
List of validated DARPin sequences.

## Author contributions

Benjamin F. N. Campbell and Shiva K. Tyagarajan conceptualized the project and designed experiments. Antje Dittmann facilitated interactome mass spectrometric analysis of gephyrin protein complexes with the Functional Genomics Center Zurich (FGCZ). Birgit Dreier and Andreas Plückthun conceptualized, designed, and supervised the *in vitro* anti-gephyrin DARPin selection and screening at the High-throughput Binder Selection facility (HT-BSF). Benjamin F. N. Campbell performed all other experiments/data analysis, and wrote the manuscript original draft. All authors contributed to manuscript writing and editing.

Name: **Benjamin F. N. Campbell**

Contribution: Conceptualisation, Methodology, Validation, Formal analysis, Investigation, Data curation, Writing – Original draft, Vizualisation, Project administration, Funding acquisition.

Competing interests: None to declare.

ORCID: 0000-0003-4814-2844

Funding: UZH Forschungskredit Candoc

Name: **Shiva K. Tyagarajan**

Contribution: Conceptualisation, writing – Original draft, Supervision, Data analysis, Project administration, Funding acquisition.

Competing interests: None To declare ORCID: 0000-0003-0074-1805

Funding: Swiss National Science Foundation (310030_192522 /1) and UZH internal funding

Name: **Antje Dittmann**

Contribution: Formal analysis, Data curation.

Competing interests: None to declare.

ORCID: 0000-0002-2570-5192

Funding: None to declare.

Name: **Birgit Dreier**

Contribution: Methodology, Resources.

Competing interests: None to declare

ORCID: None

Funding: None to declare.

Name: **Andreas Plückthun**

Contribution: Resources, Project administration, Funding acquisition.

Competing interests: A.P. is a cofounder and shareholder of Molecular Partners, who are commercializing the DARPin technology.

ORCID: 0000-0003-4191-5306

Funding: Swiss National Science Foundation (310030_192689) and UZH internal funding

## Acknowledgements

We would specifically like to thank Sven Furler, Thomas Reinberg, Joana Marinho, and Jonas Schaefer from the HT-BSF for their assistance in performing the ribosome display DARPin screen. We would like to thank Yuan-Chen Tsai and Marta Figueredo for assistance in the preparation of primary hippocampal neuron cultures. We additionally thank the Functional Genomics Centre Zurich (FGCZ) for carrying out mass-spectrometric analysis and support. We appreciated the help provided by the Protein Production and Purification Core Facility (PTPSP) Lausanne for carrying out medium-scale hFc-tagged DARPin expression. We would also like to thank the members of the Tyagarajan lab for constructive input on manuscript composition.

---

**Figure 1 Supplement 1:**
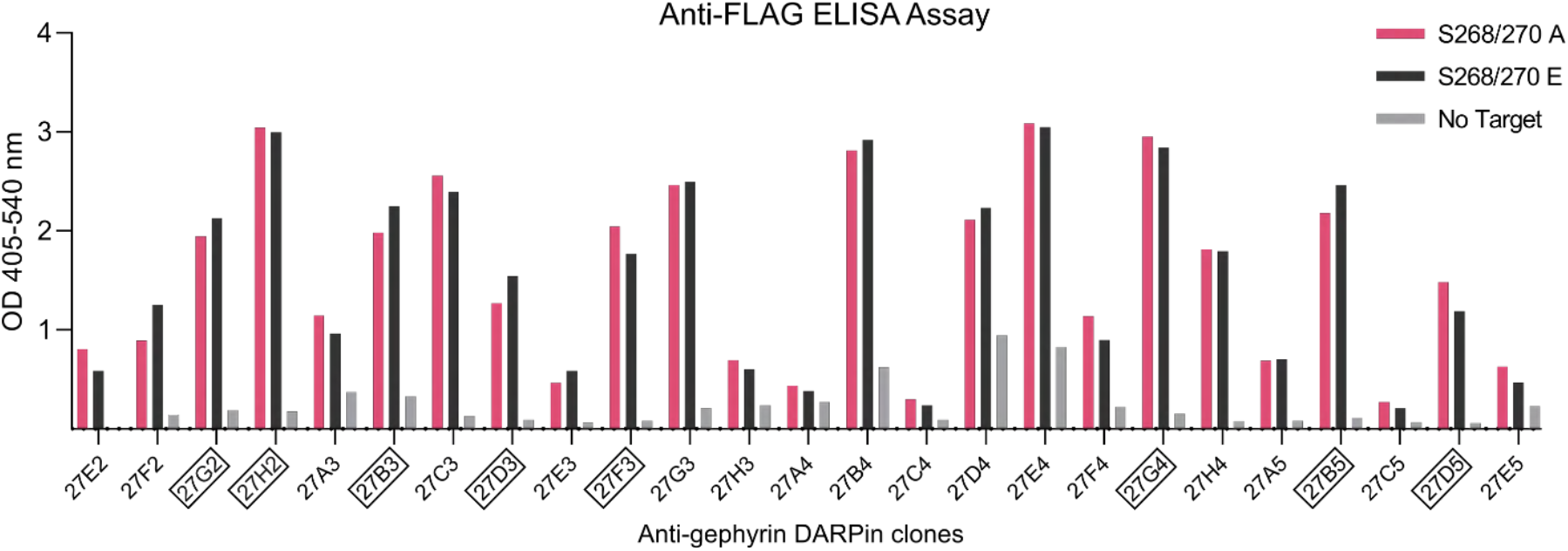
ELISA binding evaluation of anti-gephyrin DARPins. Anti-FLAG ELISA binding assay results indicating DARPin binding to phospho-null and phospho-mimetic gephyrin for 25 sequenced clones from a ribosome-display based DARPin binder selection. DARPin clones characterized further in this study are indicated in boxes.

**Figure 1 Supplement 2:**
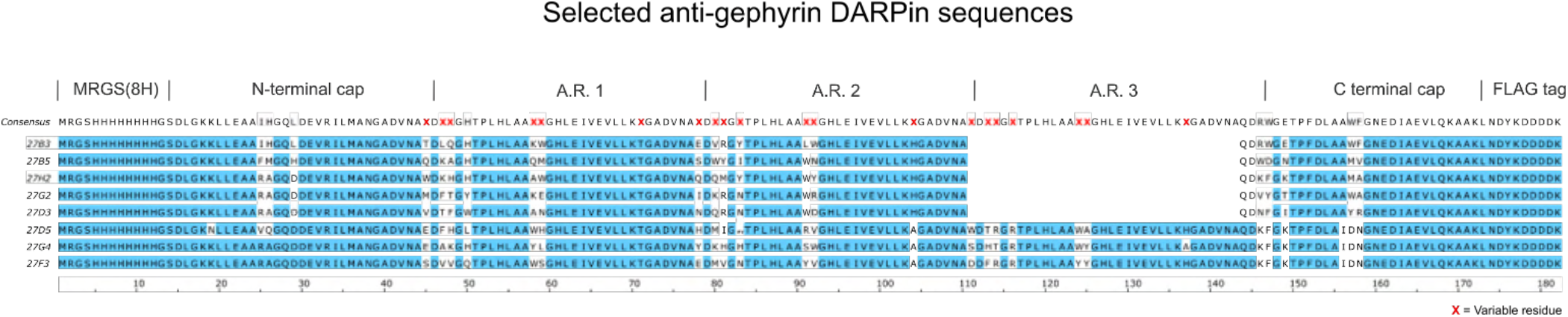
Sequence alignment of characterised anti-gephyrin DARPins. Aligned sequences of anti-gephyrin DARPins characterised in detail in this study, containing 2 or 3 randomised ankyrin repeats (A.R.). The consensus DARPin sequence is indicated above with randomised residues indicated by a red X. See materials and methods Table 6 for both DNA and protein sequences.

**Figure 2 Supplement 1.**
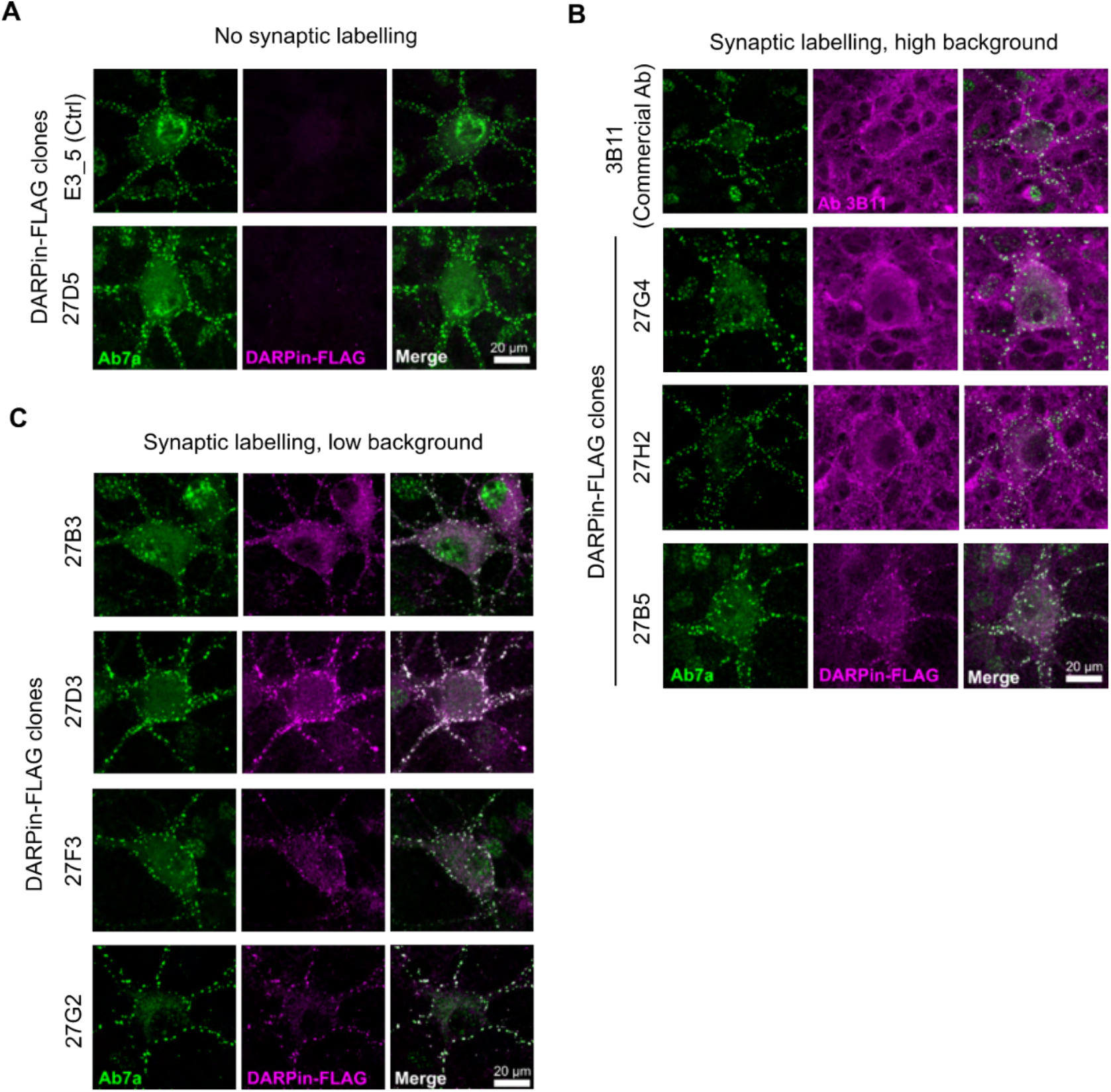
Morphological characterization of DARPin-FLAG labelling in hippocampal neuron culture. Fixed embyronic E17 rat hippocampal neuron cultures (DIV15) were stained using DARPin-FLAG clones and subsequently detected with anti-FLAG antibodies and compared to staining with commercial anti-gephyrin antibody clone Ab7a or 3B11. **A)** DARPin-FLAG control (E3_5) and clone 27D5 with no synaptic labelling. **B)** DARPin-FLAG clones and antibody 3B11 which demonstrate high background labelling. **C)** DARPin-FLAG clones with highly specific inhibitory synapse labelling.

**Figure 3 Supplement 1.**
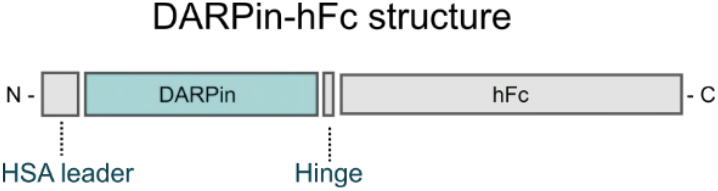
Structure of DARPin-hFc 27G2. **A**) DARPin clones were inserted into a construct containing an N-terminal HSA leader sequence for mammalian recombinant expression and a C-terminal hFc tag for detection with secondary antibodies.

**Figure 3 Supplement 2.**
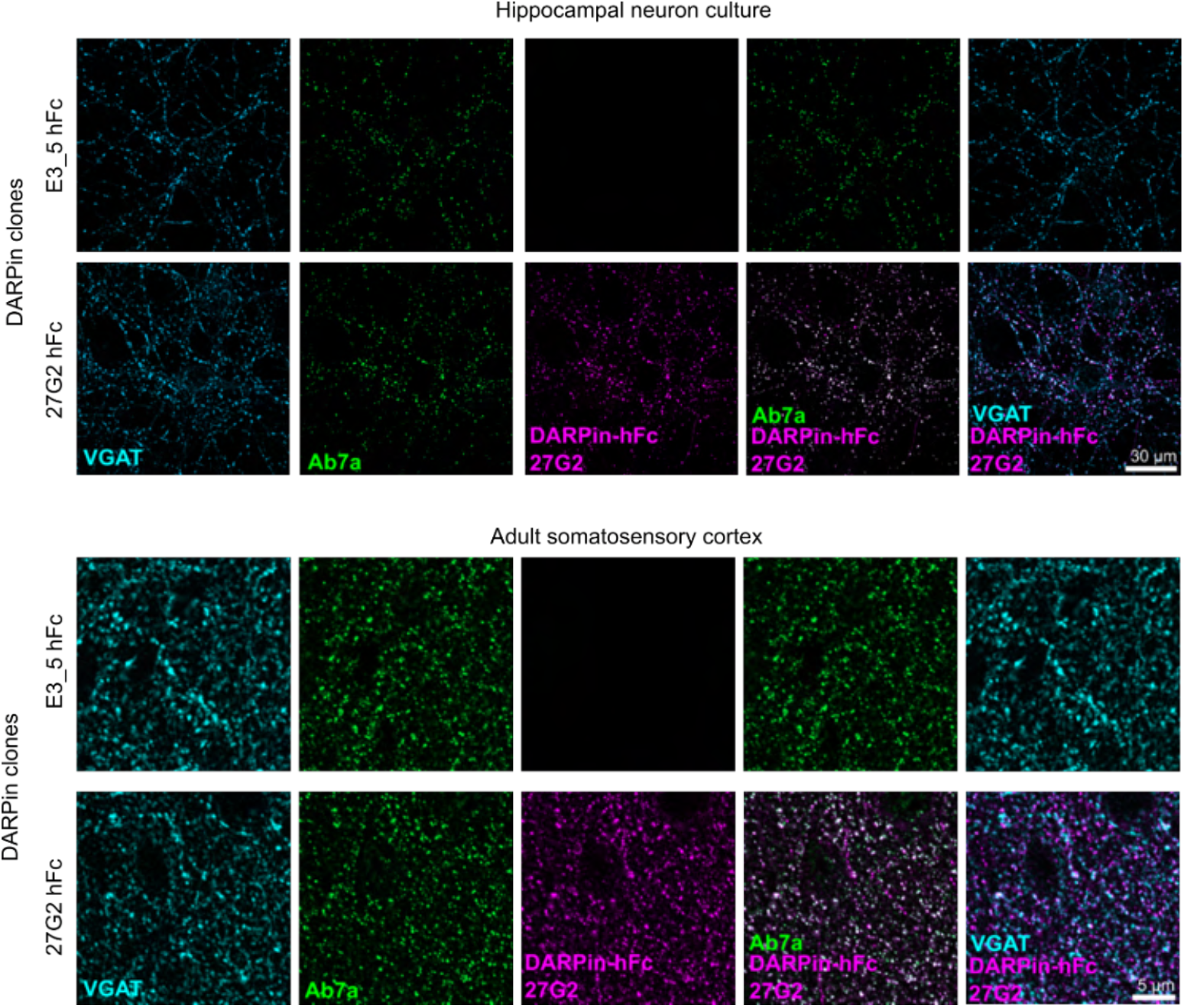
Validation of DARPin-hFc 27G2 for immunostaining. Anti-gephyrin DARPin-hFc 27G2 labels postsynaptic gephyrin puncta in hippocampal neuron culture and adult brain tissue (layer 2/3 somatosensory cortex).

**Figure 3 Supplement 3.**
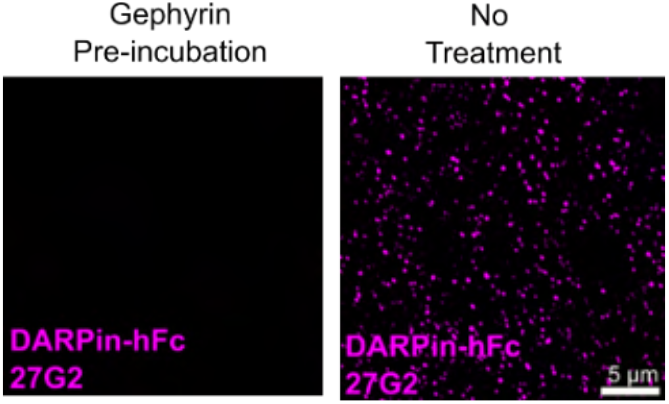
Competition with recombinant gephyrin reduces DARPin-hFc reactivity in tissue. DARPin-hFc 27G2 cluster detection is blocked by incubation with molar excess of recombinant gephyrin indicating its specificity in tissue.

**Figure 3 Supplement 4.**
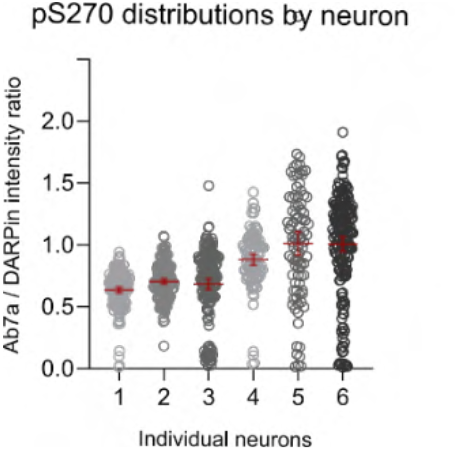
Variation in Ab7a reactivity. The ratio of fluorescent intensity signal between pS270-specific antibody Ab7a and the phosphorylation non-specific DARPin-hFc 27G2 indicates that Ab7a labelling is variable between clusters within and between individual synapses and neurons. Each data represents one cluster analysed from 6 individual example neurons with different patterns of relative Ab7a reactivity. Median and SD are indicated in red. Figure 3 – Source Data 2: contains the values used to plot Figure 3. Supplement 4.

**Figure 4 Supplement 1:**
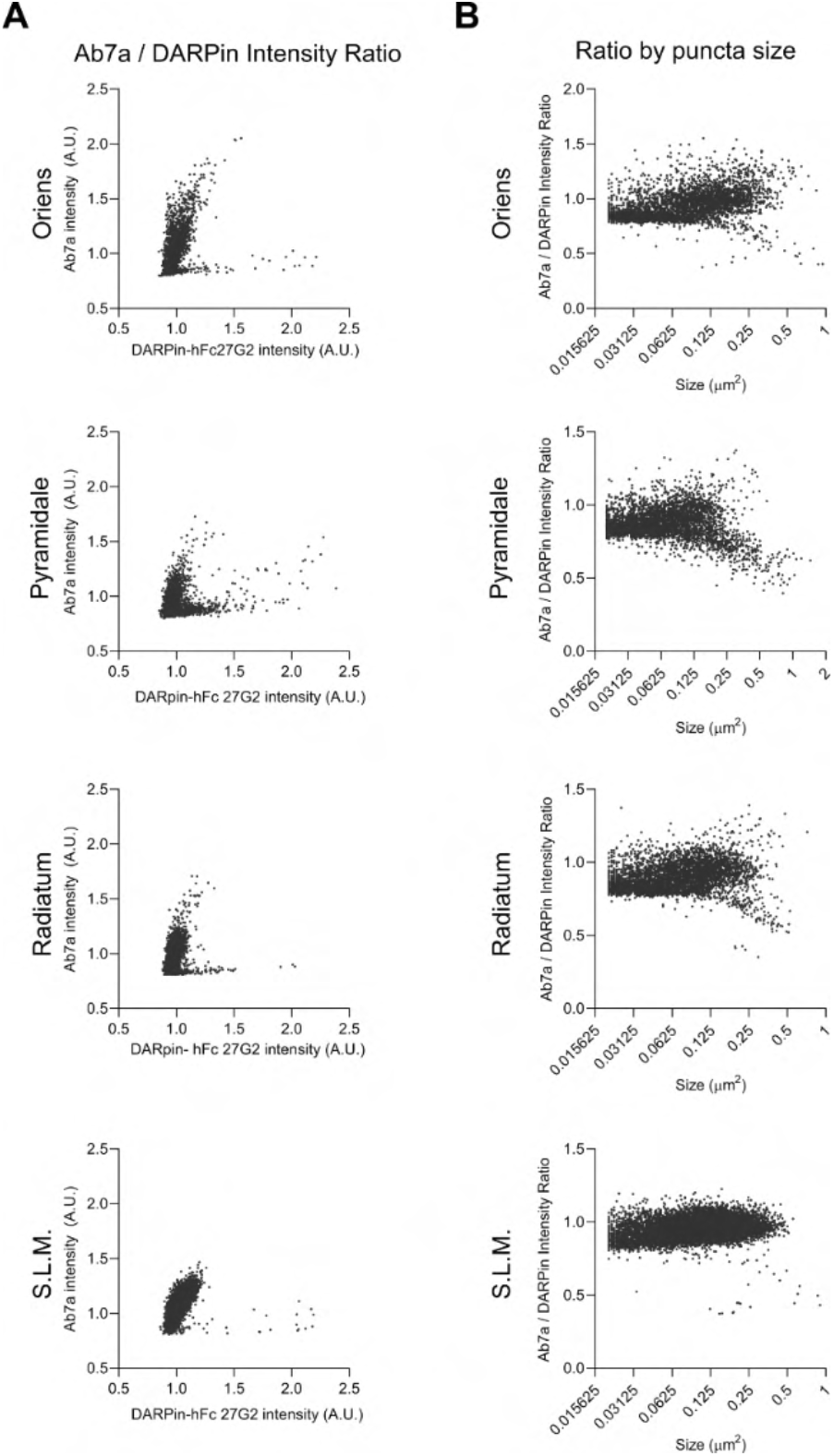
Relative pS270 synaptic distribution in the hippocampal CA1. Extended example distribution of signal from adult brain tissue from Figure 4. including the s. oriens, pyramidale, radiatum, and stratum lacunosum moleculare (S.L.M.) **A)** Ab7a versus DARPin-hFc 27G2 puncta intensity. **B)** Ab7a/DARPin-hFc 27G2 intensity ratio plotted by puncta size. Figure 4 – Source Data 2: Contains data and statistical analysis presented in Figure 4 Supplement 1 A and B.

**Figure 5 Supplement 1.**
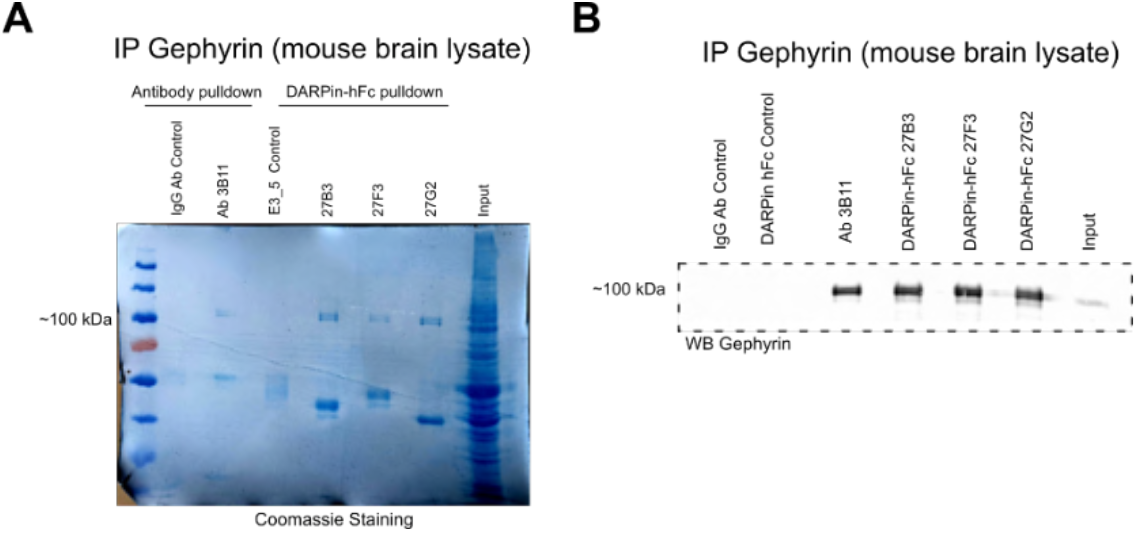
Anti-gephyrin DARPins affinity purify gephyrin from mouse brain lysates. **A)** Coomassie stained acrylamide gel indicating abundant gephyrin precipitated both by the antibody 3B11 and DARPin-hFc 27B3, 27F3, and 27G2 without signal in antibody (IgG) or DARPin (E3_5) controls. Lower bands correspond to IgG or DARPin-hFc protein. **B)** Immunoblot of gephyrin precipitated with different binders probed with the antibody 3B11. Figure 5 – Source Data 2: Raw Coomassie gel images and immunoblots from Figure 5 Supplement 1.

**Figure 5 Supplement 2.**
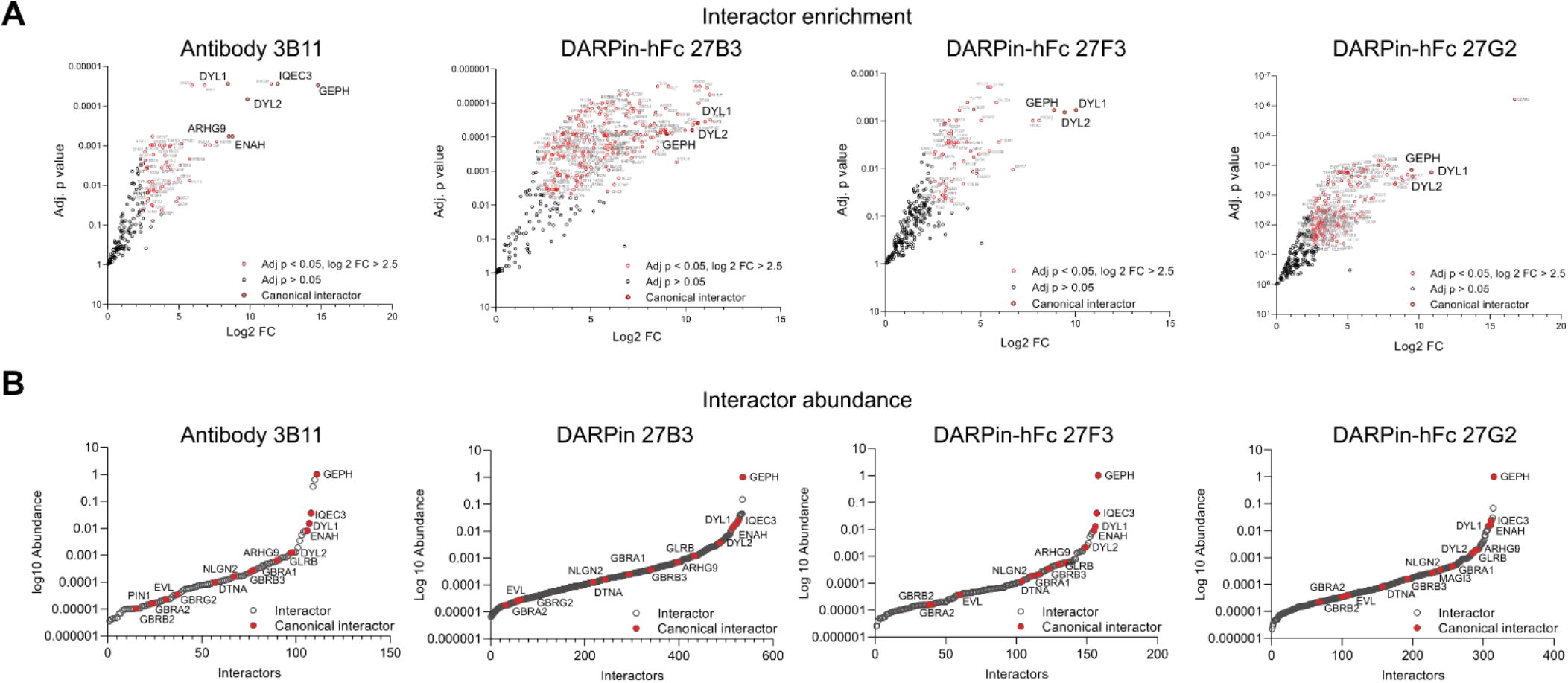
Interactor identification plots. **A)** Volcano plots of enriched proteins with the Log_2_ FC >2.5 and FDR-adjusted p-value compared to controls. Red points indicate identified gephyrin interacting proteins, with canonical interactors indicated by enlarged text. **B)** Abundance of gephyrin interactors for antibody and DARPin-hFc experiments with canonical interactors indicated in red demonstrating several orders of magnitude difference. interactors. Figure 5 – Source Data 3: Identity and quantification of abundance of interacting proteins presented in Figure 5 Supplement 2. Figure 5 – Source Data 4: Compiled list of proteins from all gephyrin interactor experiments used to assess gephyrin interactor identity.

**Figure 5 Supplement 3.**
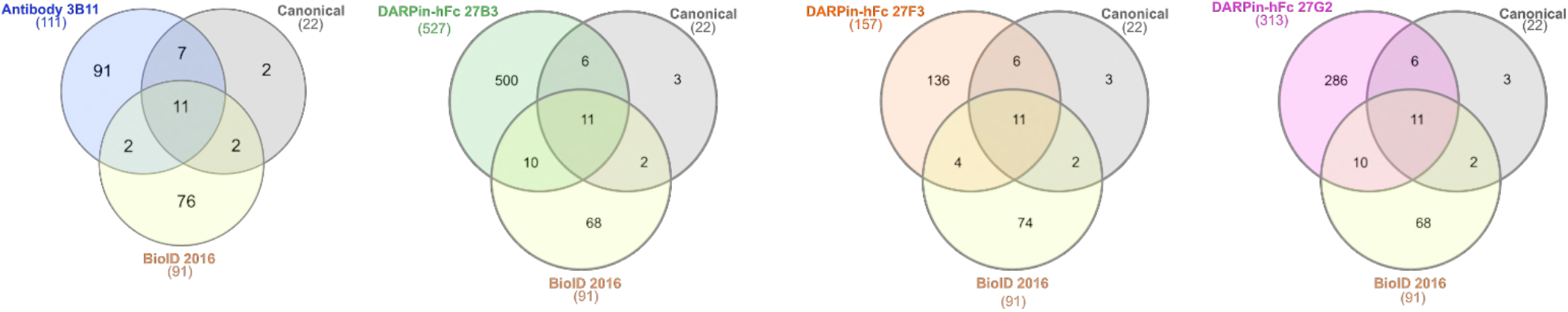
Interactome overlap with previous literature. Venn diagrams showing the overlap in identified interactors determined using both antibody and DARPin-based interactomes compared to previously identified interactors from the literature (see methods) and by and BioID (Uezu et al., 2016).

**Figure 5 Supplement 4.**
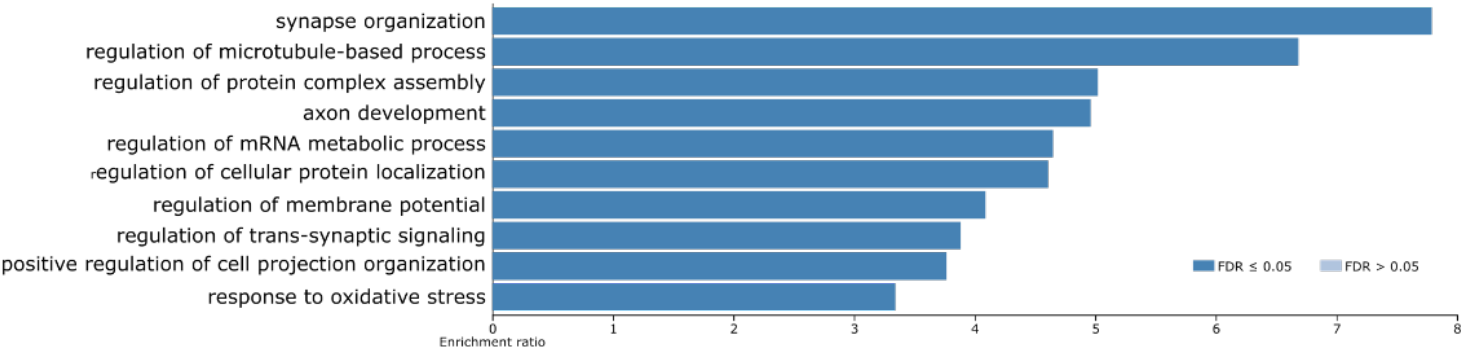
Ontological enrichment analysis of the consensus gephyrin interactome. Biological process enrichment (WebGestalt) for the 120 consensus gephyrin interactors showing significantly regulated ontologies.

**Figure 6 Supplement 1.**
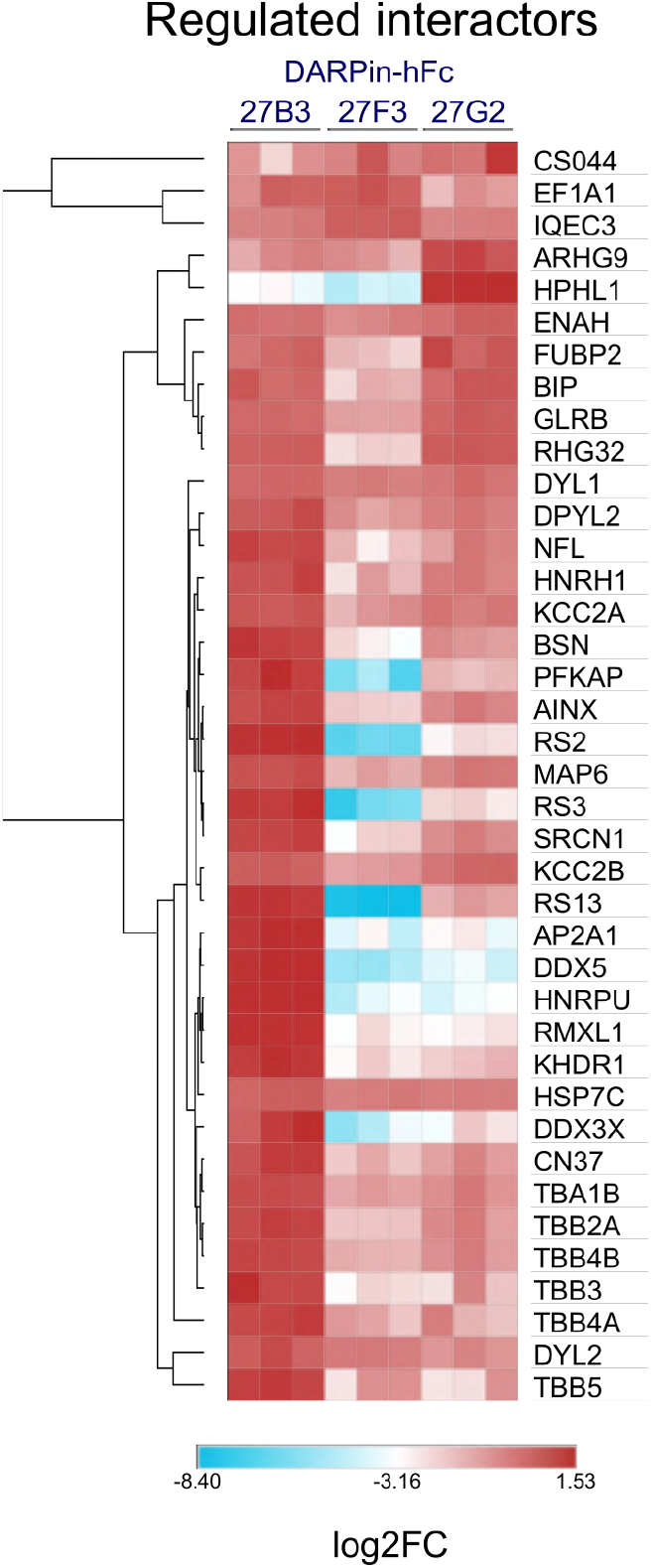
DARPin-specific gephyrin interactor abundance. **A)** Common gephyrin interactors identified by all DARPin-hFc-based interactomes showing proteins with significantly different abundances relative to gephyrin, organised by hierarchical clustering. Only significantly regulated interactors are shown. **Statistics:** Two-way ANOVA with multiple comparisons correction comparisons all groups, 3 replicates per group. Figure 6 – Source Data 2: Values and statistical test results indicating differentially abundant gephyrin interactors between binding experiments.

**Figure 6 Supplement 2.**
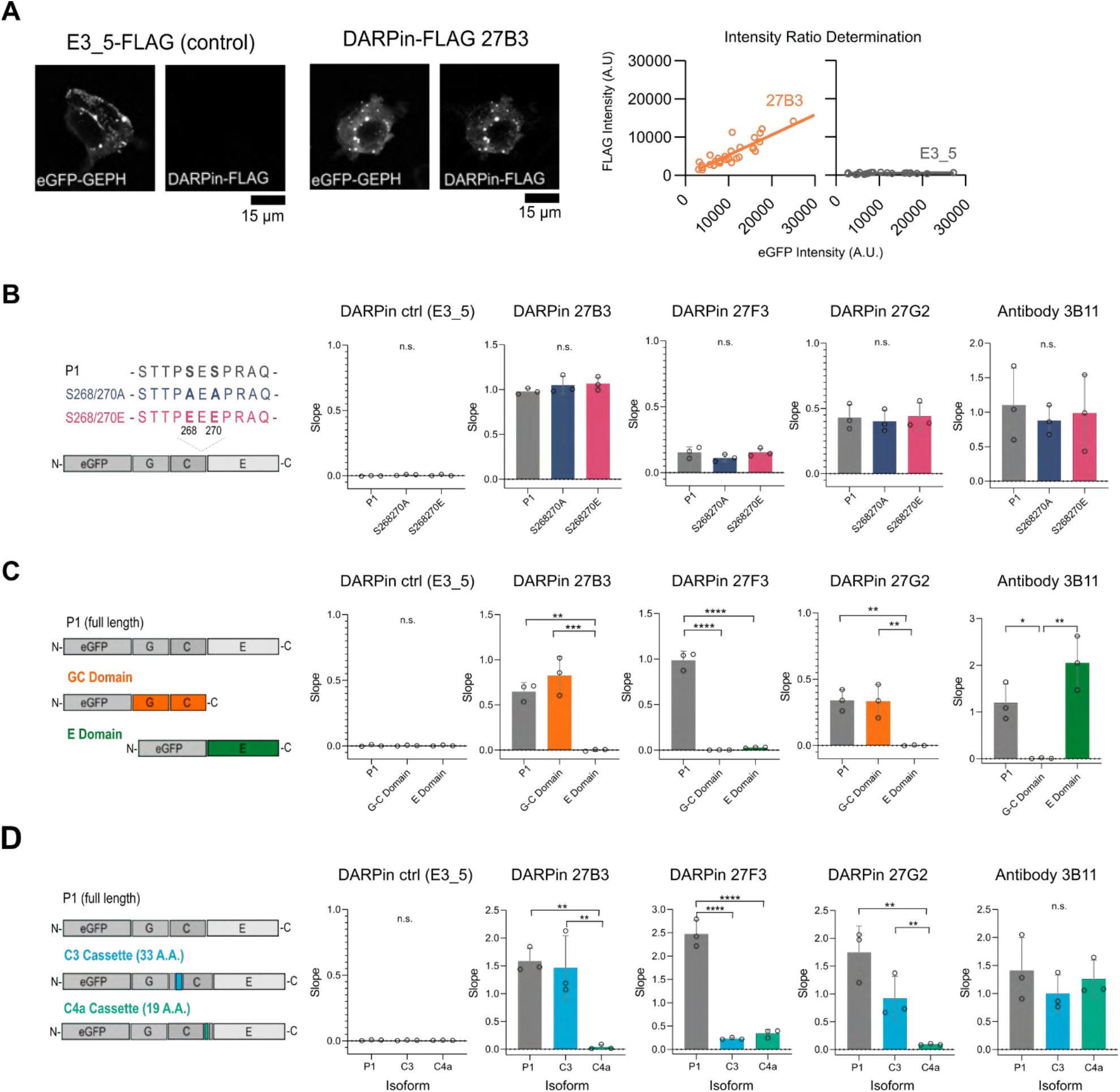
Identification of gephyrin-binding preferences of anti-gephyrin DARPins using an in-cell HEK293T fluorescence assay. **A)** Representative images of eGFP-gephyrin expressed in HEK cells which were fixed and probed using DARPin-FLAG clones or commercial antibody clone 3B11. Shown is eGFP and FLAG signal provided by the control (E3_5) and gephyrin-binding DARPin-FLAG clones (e.g. 27B3). the relative signal between eGFP and FLAG for a given cell are plotted, and the slope compared between clones to assess relative binding. **B)** Quantification of binder labelling of eGFP-tagged gephyrin WT versus S268A/S270A and S268E/S270E phospho-mutants overexpressed in HEK293T cells. **C)** Quantification of binding to overexpressed full length (P1 variant) gephyrin or GC or E domains only. **D)** Quantification of binding to eGFP-tagged gephyrin P1 isoform or isoforms including the C3 or C4a cassettes. **Statistics:** One-way ANOVA, * p<0.05, ** p<0.01, *** p<0.001, **** p<0.0001. Data points represent the slope calculated from at least 25 cells in 3 independent experiments. All panels: mean and SD are presented. Figure 6 – Source Data 3: Values and statistical analysis performed to generate graphs in Figure 6 Supplement 2 panels B, C, and D.

**Figure 6 Supplement 3.**
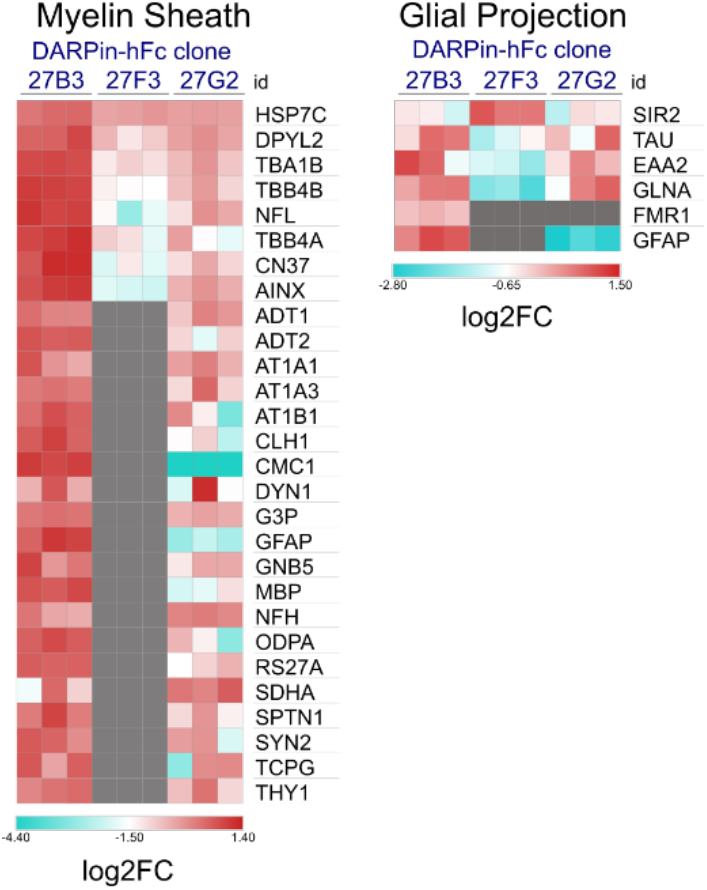
Non-neuronal interactor ontology. Heatmap of relative abundance of proteins of “myelin sheath” or “glial projection” ontology between different DARPin-detected interactomes, grey squares indicate that the binder was not detected as a gephyrin interactor using a given DARPin. Figure 6 – Source Data 4: Values used to generate heat maps in Figure 6 Supplement 3.

